# *RGA1* alleviates low-light-repressed pollen tube growth by improving the metabolism and allocation of sugars and energy

**DOI:** 10.1101/2022.09.25.509406

**Authors:** Hubo Li (李沪波), Baohua Feng (奉保华), Juncai Li (李俊材), Weimeng Fu (符卫蒙), Wenting Wang (王文婷), Tingting Chen (陈婷婷), Lianmeng Liu (刘连盟), Zhihai Wu (武志海), Shaobing Peng (彭少兵), Longxing Tao (陶龙兴), Guanfu Fu (符冠富)

## Abstract

Low-light stress compromises photosynthetic and energy efficiency and leads to spikelet sterility; however, the effect of low-light stress on pollen tube elongation in the pistil remains poorly understood. The gene *RGA1*, which encodes a Gα subunit of the heterotrimeric G protein, enhanced low-light tolerance in rice plants at anthesis by preventing the cessation of pollen tube elongation in the pistil. The levels of reactive oxygen species were higher and the content of ATP and ATPase was lower in *RGA1* mutant (d1) plants compared with wild-type and *RGA1*-overexpressing (OE-1) plants under low-light conditions. Energy deficits, rather than interference with signaling transduction pathways, were the main contributors to the inhibition of pollen tube elongation in the pistil by low-light stress. In this process, marked increases in the activities of acid invertase (INV), sucrose synthase (SUS), and mitochondrial respiratory electron transport chain complexes, as well as the relative expression levels of *SUTs*, *SWEETs*, *SUSs*, *INVs*, *CINs*, *SnRK1A*, and *SnRk1B*, were observed in OE-1 plants. INV and ATPase activators (sucrose and Na_2_SO_3_, respectively) increased spikelet fertility by improving the energy status in the pistil under low-light conditions, and the ATPase inhibitor Na_2_VO_4_ induced spikelet sterility and decreased ATPase activity. Therefore, *RGA1* could alleviate the low-light stress-induced impairment of pollen tube elongation to increase spikelet fertility by promoting sucrose unloading in the pistil and improving the metabolism and allocation of energy.

## Introduction

Rice (*Oryza sativa*) is a critically important crop that feeds more than half of the world’s population (Horie, 2019). However, increases in yield are impeded by adverse environmental factors, such as extreme temperatures, drought, salt stress, and low-light stress, which might increase in severity with continued global warming driven by industrial development and anthropogenic activities (Horie, 2019; Long et al., 2022). China has faced various challenges associated with dimming, which is associated with alterations in the amount of solar radiation reaching the Earth’s surface due to long-term continuous rain and increases in aerosol pollution, and these problems have only been exacerbated by rapid urbanization and economic development (Tollenaar et al., 2017; Shao et al., 2021). Rice yield has decreased by 30–50% in some areas that experience cloudy and rainy conditions for long periods (Li et al., 2019; Kumar et al., 2020). Furthermore, approximately 0.4% of global average crop yields have been lost due to global dimming (Proctor, 2021). Therefore, low light levels have become a major threat to food security worldwide (Wang et al., 2015).

Plants use light not only as an energy source for photosynthesis but also as an environmental signal for plant growth and development (Gommers et al., 2013; Schwenkert et al., 2022). The intensity, wavelength, and direction of light have direct effects on plant morphology, physiology, and biochemistry (Fukuda et al., 2008; Shao et al., 2020; Fernández-Milmanda & Ballaré, 2021). Solar energy deficits disturb the growth and development of rice plants and result in reduced dry matter weight accumulation, yield, and quality of rice, especially at the reproductive stage (Deng et al., 2018; Ganguly et al., 2020; Deng et al., 2021). Spikelet fertility is one of the most important factors affecting rice yield, and it is determined by pollen viability, anther dehiscence, the number of pollen grains on the stigma, pollen germination, and pollen tube elongation in the pistil; it is particularly important under abiotic stress (Xu et al., 2017; Xiang et al., 2019; Jagadish, 2020; Deng et al., 2021). A large decrease in the seed-setting rate and filled grains per panicle has been observed in plants under low-light conditions at anthesis (Deng et al., 2021). This might be mainly ascribed to the lower fertilization rate, which inhibits key pollination and fertilization processes, especially anther dehiscence and pollen grain release (Deng et al., 2021). Pollen tube elongation in the pistil also regulates spikelet fertility under abiotic stresses such as heat stress (Fu et al., 2016; Zhang et al., 2018a). However, the response of pollen tube elongation to low-light stress has not yet been documented in rice.

The pollen tube is fundamental for the reproduction of seed plants (Ge et al., 2017; Yu et al., 2021; Zhong et al., 2022). The pollen tube grows quickly, and it is the channel through which sperm cells of the male gametophyte reach the ovule of the female gametophyte (Hoffmann et al., 2020). In this process, pollen tube integrity and burst are controlled by the binding of RALF family peptides to CrRLK1L family receptors and cell wall leucine-rich repeat extensins (Ge et al., 2017; Li & Yang, 2018). The pollen tube comprises highly polarized tip-growing cells and requires the coordination of cell structure and function, including intracellular turgor pressure, ion flow, skeletal microfilament dynamics, vesicle transport, entosis, exocytosis, and cell wall construction (Rottmann et al., 2016). Such energy-demanding processes depend on cytosolic pH gradients for signaling and growth (Goetz et al., 2017; Ge et al, 2019), and the autoinhibited plasma membrane proton (H^+^) ATPases act as energy transducers to sustain the ionic circuit that defines the spatial and temporal profiles of cytosolic pH (Hoffmann et al., 2020). This controls the downstream pH-dependent mechanisms essential for pollen tube elongation in the pistil. Thus, energy deficits inhibit pollen tube elongation in the pistil and result in spikelet sterility under heat stress (Zhang et al., 2018a; Jiang et al., 2020). Higher invertase activity can alleviate the impairment of pollen tube elongation under heat stress in rice plants, which enhances sucrose metabolism to maintain energy homeostasis (Jiang et al., 2020). Furthermore, plants with strong abiotic resistance show high NADH dehydrogenase and ATPase activity and thus greater energy production efficiency and utilization (De Block & Van Lijsebettens, 2011; Yu et al., 2020). Low light levels inhibit photosynthesis but have little effect on respiration; this eventually leads to carbon starvation. Insufficient carbon assimilation caused by shading markedly increases the number of empty spikelets by increasing rice spikelet sterility (Kobata et al., 2013). Therefore, low-light stress might reduce rice fertilization by affecting energy acquisition in the stigma.

Recently, we identified a low-light-intolerant mutant of *RGA1*, which encodes the Gα subunit of the heterotrimeric G protein (d1) in rice (Ferrero-Serrano et al., 2018). It is a signal transduction protein involved in various abiotic stress responses, including the response to heat stress, cold stress, drought stress, and salt stress (Jangam et al., 2016). Heat stress tolerance is high in tobacco plants overexpressing *RGA1* (Misra et al., 2007). However, *RGA1* mutant plants show greater drought tolerance, as well as higher stomatal conductance, lower leaf temperatures, and delayed onset of the stomatal and non-stomatal limitation of photosynthesis (Ferrero-Serrano & Assmann, 2016). *RGA1* mutant plants also show salt tolerance, photoavoidance, and photoprotection (Urano et al., 2014; Colaneri et al., 2014; Ferrero-Serrano et al., 2018). However, the response of *RGA1* to low-light conditions has not yet been documented in rice plants. We found that *RGA1* can enhance spikelet fertility by preventing the cessation of pollen tube growth in the pistil under low-light stress. In this process, activities of sucrose synthase (SUS), acid invertase (INV), ATP and ATPase, as well as the relative expression levels of sucrose transporters (*SUTs*), sugars will eventually be exported transporters (*SWEETs)*, and sucrose-non-fermenting 1-related kinase 1 (*SnRK1s*), were higher in plants overexpressing *RGA1* than those of its mutant plants under low-light conditions. We hypothesized that *RGA1* might improve sucrose and energy metabolism and allocation in the pistil to enhance pollen tube elongation under low-light stress. We characterized pollen tube morphology and measured levels of carbohydrates, SUS and INV activity, sugar and energy metabolism, and antioxidant capacity to clarify the mechanism by which *RGA1* confers low-light tolerance in rice plants.

## Results

### *RGA1* enhances dry matter weight, grain yield, yield components, and net photosynthesis under low-light conditions

The morphology of plants and the panicle varied among the three rice genotypes under low-light conditions (Fig. 1, a–d). Total dry matter weight, including the sheath, stem, leaf, and panicle, was decreased by low-light stress, and reductions of 19.9%, 37.5%, and 13.9% were observed in WT, d1, and OE-1 plants, respectively (Fig. 1e). Similar patterns were observed in the ratio of panicle dry matter weight to total dry matter weight; specifically, low-light stress resulted in reductions of approximately 30.7%, 67.2%, and 7.2% in WT, d1, and OE-1 plants, respectively (Fig. S1). Marked reductions in grain yield of 44.3%, 84.7%, and 20.7% were observed in WT, d1, and OE-1 plants under low-light conditions compared with their respective controls, respectively; reductions in spikelet fertility of approximately 44.7%, 91.6%, and 14.1% were observed in WT, d1, and OE-1 plants compared with their respective controls, respectively (Fig. 1, f and g). No difference in kernel weight was observed between control and low-light conditions, and the greatest decrease was observed in d1 plants under low-light conditions (Fig. 1h). No differences in net photosynthetic rate were observed among these three genotypes under both control and low-light conditions, but the net photosynthetic rate was slightly higher in d1 plants than in WT and OE-1 plants (Fig. S2).

**Figure 1.**
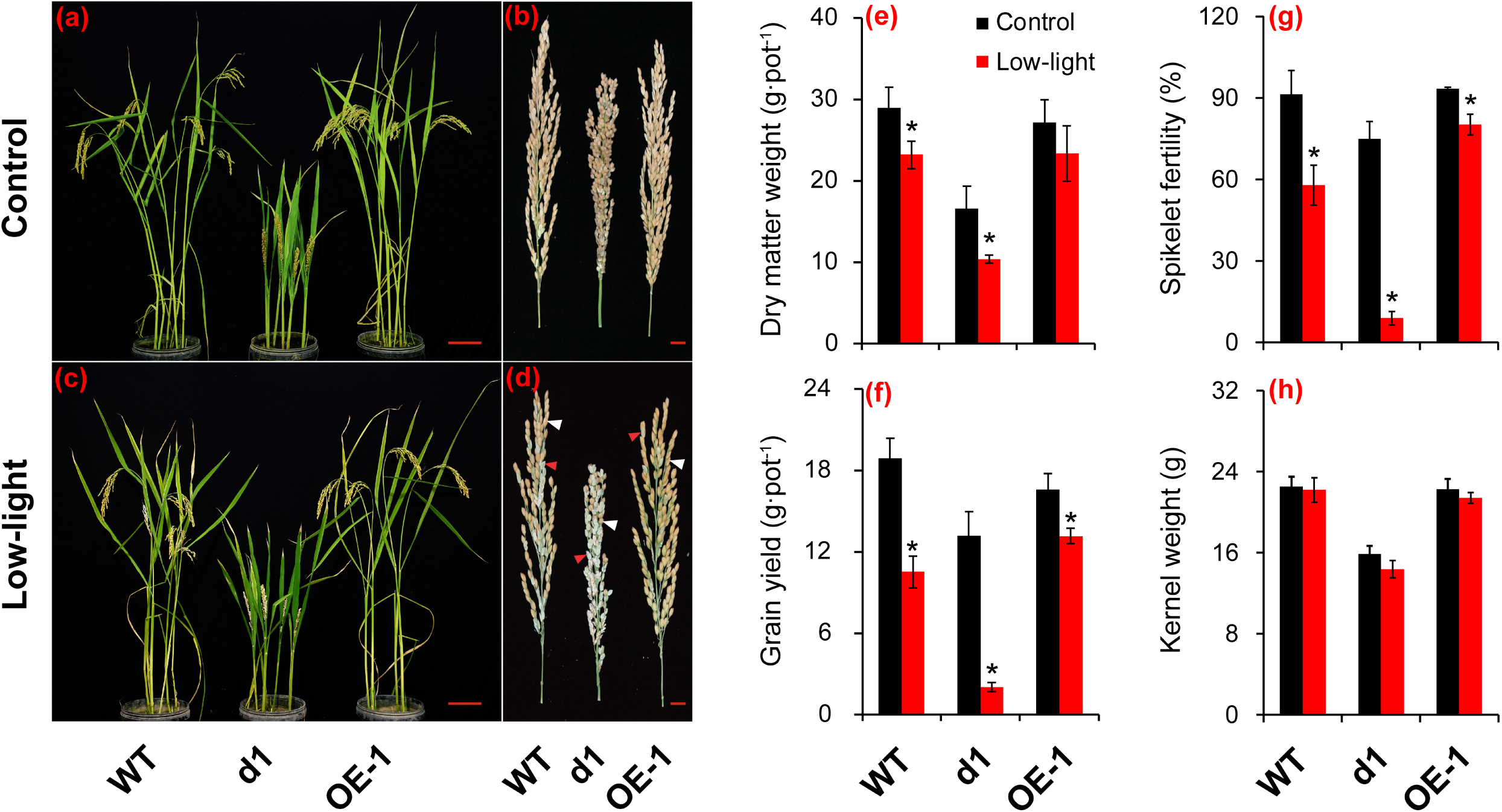
Effect of low-light stress at anthesis on the dry matter weight, yield, and yield components of rice. **(a, c)**, The morphology of rice plants. Bar = 5 cm; **(b, d)**, Morphology of the panicle. Bar = 1 cm; **e**, Dry matter weight of plants in each pot; **f**, Grain yield of plants in each pot; **g**, Spikelet fertility; **h**, Kernel weight. The mean values and standard errors in the figures correspond to data from five replicates. Student’s *t*-tests were conducted to evaluate the significance of differences between treatments within single genotypes. * indicates P-value < 0.05. Red triangles indicate sterile grains, and white triangles indicate fertile grains.

### *RGA1* prevents the low-light stress-induced cessation of pollen tube growth in the pistil

Spikelet fertility was the main factor contributing to differences in grain yields among the three genotypes, and pollen viability, anther dehiscence, number of pollen grains on the stigma, and pollen tube elongation in the pistil were studied. There were no differences in pollen viability, anther dehiscence, number of pollen grains on the stigma, and pollen germination rate between control and low-light conditions (Fig. S3; Fig. S4; Fig. 2A, g and h). Pollen tube elongation was inhibited by low-light stress (Fig. 2A, a–f and g). The proportion of pollen tubes that penetrated the ovule was approximately 59.8%, 51.8%, and 64.2% in WT, d1, and OE-1 plants under normal temperature conditions, respectively; however, the proportion of pollen tubes that penetrated the ovule was 24.6%, 7.5%, and 40.0% lower under low-light conditions in WT, d1, and OE-1 plants, respectively.

**Figure 2.**
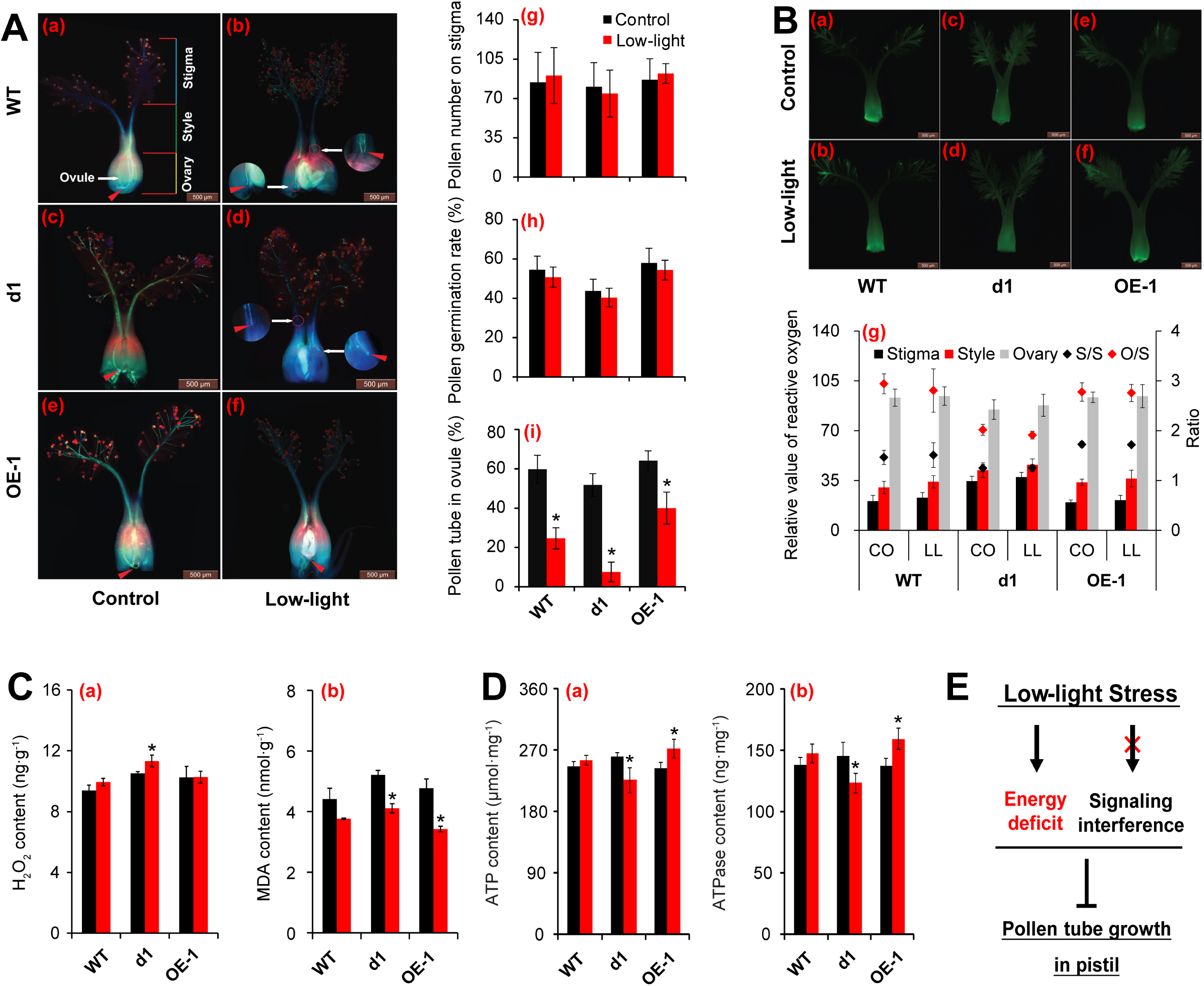
Effect of pollen tube elongation and changes in energy status and ROS signaling in the pistil of rice under low-light stress at anthesis. **A (a-f)**, The morphology of pollen tube elongation in pistil. The triangles indicate pollen tube elongation in pistil, while the circle indicates the cease of pollen tube. Bar = 500 μm. **(g-i)**, Changes in pollen viability, pollen numbers and pollen germination on stigma and pollen tube elongation in pistil under low-light condition. **g**, Pollen numbers on stigma; **h**, Pollen germination on stigma; **i**, Pollen numbers in ovule. The mean values and standard errors in the figures correspond to data from ten replicates. **B (a-f)**, ROS fluorescence staining of pistil. Bar = 500 μm; **g**, Statistics of ROS fluorescence intensity of style, stigma and ovary of pistil and the ratio of style/stigma and ovary/style. The mean values and standard errors in the figures correspond to data from five replicates. **C**, Effect of low-light stress on H_2_O_2_ and MDA contents of pistil of rice. The mean values and standard errors in the figures correspond to data from three replicates. **D**, Effect of low-light stress on ATP and ATPase contents of pistil of rice. The mean values and standard errors in the figures correspond to data from three replicates. **E**, Schematic diagram of the mechanism of low-light stress inhibiting pollen tube elongation. Student’s *t*-tests were conducted to evaluate the significance of differences between treatments within single genotypes. * indicates P-value < 0.05.

### Energy deficits inhibit pollen tube elongation rather than signaling interference under low-light stress

ROS are important signaling molecules that play a role in guiding the elongation of the pollen tube into the ovule. Thus, the distribution of ROS in the pistil was characterized, and the H_2_O_2_ level was determined (Fig. 2B and C). The ROS fluorescence images of pistils under low-light stress revealed that the fluorescence intensity was slightly higher in the three genotypes under control conditions compared with low-light conditions (Fig. 2B, a–f). The relative values of ROS of the style, stigma, and ovary of the three genotypes were higher under low-light conditions compared with control conditions, but none of these differences were significant (Fig. 2B, g). There were also no significant differences in the ratio of the relative values of ROS of the style/stigma and ovary/style of the three genotypes under the two treatments, indicating that low-light stress had no effect on the transduction of ROS signals (Fig. 2B, g). The H_2_O_2_ level was higher in all three genotypes under low-light conditions compared with their respective controls, and the greatest increase was observed in d1 plants (Fig. 2C, a). The MDA content was lower in these three genotypes under low-light conditions than under control conditions (Fig. 2C, b).

Pollen tube elongation is an energy-demanding process; thus, the energy status of seed plants, as well as the execution of signaling transduction pathways, is important for successful pollination. The content of ATP and ATPase varied among the three rice genotypes under low-light conditions (Fig. 2D, a and b). No differences in the content of ATP and ATPase were observed in WT plants between control and low-light conditions; however, the content of ATP and ATPase was lower in d1 plants under low-light stress than under control conditions. By contrast, the content of ATP and ATPase in OE-1 plants was significantly higher under low-light conditions than under control conditions.

### The function of *RGA1* in sugar transport in the pistil under low-light conditions

There was no noticeable difference in the content of soluble sugar, starch, and NSC in the three genotypes between control and low-light conditions (Fig. 3A, a, b and c). The content of soluble sugar, starch, and NSC was slightly higher in the three rice genotypes under low-light conditions compared with control conditions, with the exception of the starch content in the pistil of d1 plants; the greatest increases in the content of soluble sugar, starch, and NSC between low-light and control conditions were observed in OE-1 plants, followed by WT and d1 plants.

**Figure 3.**
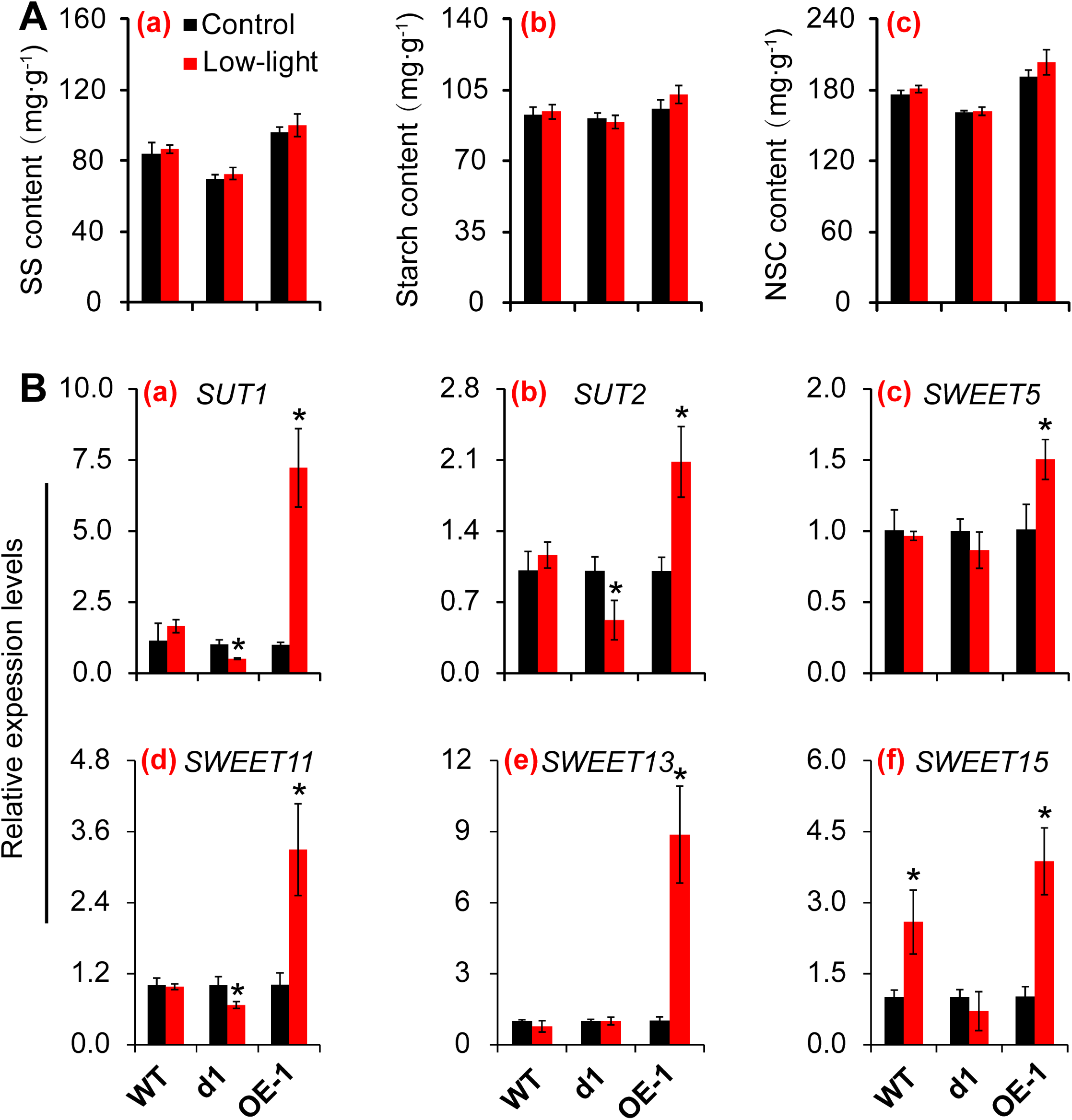
Effect of low-light stress on sugar transport in the pistil of rice at anthesis. Effect of low-light stress on sugar transport in pistil of rice at anthesis. **A**, Changes in contents of soluble sugar, starch and NSC in pistil of rice under low-light condition. **a**, Soluble sugar; **b**, Starch; **c**, NSC. **B**, Changes in expressions of *SUTs* and *SWEETs* genes in pistil of rice under low-light condition. The mean values and standard errors in the figures correspond to data from three replicates. NSC, Non-structural carbohydrates; *SUTs*, Sucrose transporters; *SWEETs*, Sugars will eventually be exported transporters. Student’s *t*-tests were conducted to evaluate the significance of differences between treatments within single genotypes. * indicates P-value < 0.05.

SUTs and SWEETs are involved in the allocation of assimilates in plants, and several genes involved in sucrose transport were studied. The expression patterns of *SUT1* and *SUT2* were similar in the three genotypes. There were no differences in the relative expression levels of *SUT1* and *SUT2* in WT plants between control and low-light conditions (Fig. 3B, a and b). In d1 plants, the expression of these two genes was significantly inhibited by low-light stress; however, the expression of *SUT1* and *SUT2* was higher in OE-1 plants under low-light conditions compared with control conditions. There were no differences in the expression of *SWEET5* and *SWEET13* in WT and d1 plants between control and low-light conditions; however, the expression of these two genes was higher in OE-1 plants under low-light conditions compared with control conditions (Fig. 3B, c and e). Variation in the expression patterns of *SWEET11* and *SWEET15* were observed among the three rice genotypes (Fig. 3B, d and f). Low-light stress had little effect on the expression of *SWEET11* in WT plants; however, low-light stress strongly inhibited the expression of this gene in d1 plants. Low-light stress greatly increased the expression of *SWEETs* in OE-1 plants. Low-light stress increased the expression of *SWEET15* in WT and OE-1 plants, and the magnitude of the increase was higher in OE-1 plants than in WT plants (Fig. 3B, d). Low-light stress had little effect on the expression of *SWEET15* in d1 plants (Fig. 3B, f).

### *RGA1* promotes sucrose metabolism in the pistil under low-light conditions

There were no differences in the content of sucrose and fructose in WT and OE-1 plants between control and low-light conditions, and the content of sucrose was significantly higher in d1 plants under low-light conditions than under control conditions (Fig. 4A, a and c). The content of glucose was higher in OE-1 plants under low-light conditions than under control conditions; however, no difference in the glucose content was observed in WT and d1 plants between low-light and control conditions (Fig. 4A, b). The ratio of sucrose to soluble sugar, which includes sucrose, glucose, and fructose, varied little in WT and OE-1 plants under low-light conditions; the ratio of sucrose to soluble sugar was higher in d1 plants under low-light conditions by comparison (Fig. 4A, d).

**Figure 4.**
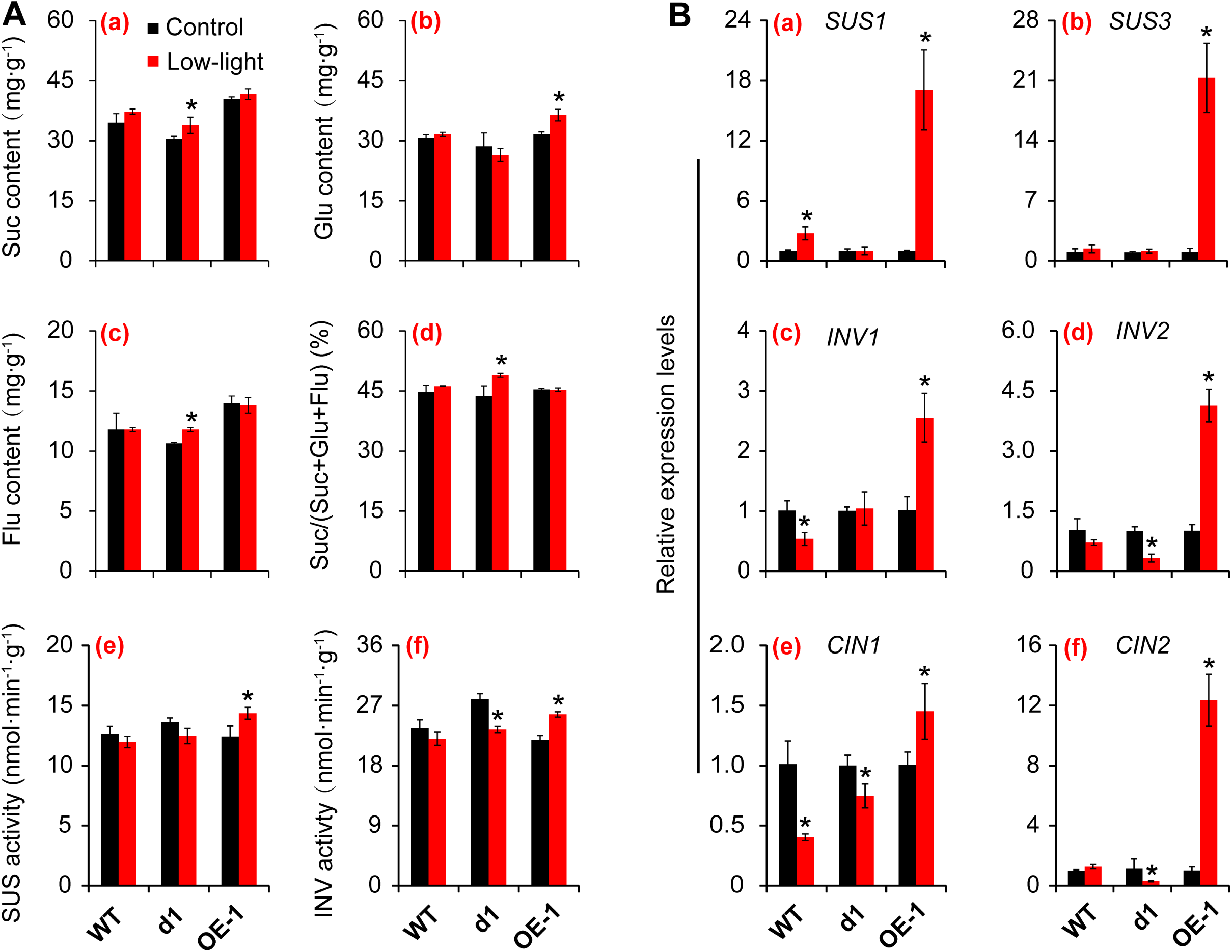
Effect of low-light stress on sucrose metabolism in the pistil of rice at anthesis. **A**, Changes in sugar contents as well as activities of SUS and INV in pistil of rice under low-light condition. **a**, Sucrose content; **b**, Glucose content; **c**, Fructose content; **d**, Ratio of sucrose to total soluble sugar including sucrose, glucose and fructose; **e**, SUS activity; **f**, INV activity. **B**, Expressions of genes related to SUS and INV in pistil under low-light condition. **a**, *SUS1*; **b**, *SUS3*; **c**, *INV1*; **d**, *INV2*; **e**, *CIN1*; **f**, *CIN2*. The mean values and standard errors in the figures correspond to data from three replicates. Student’s *t*-tests were conducted to evaluate the significance of differences between treatments within single genotypes. * indicates P-value < 0.05.

SUS and INV play key roles in sucrose metabolism. Slight reductions in SUS activity were observed in WT and d1 plants under low-light conditions compared with control conditions, but these differences were not significant (Fig. 4A, e). In OE-1 plants, low-light stress increased SUS activity. Patterns of INV activity varied among these three genotypes (Fig. 4A, f). There was no difference in INV activity in WT plants between control and low-light conditions. INV activity was markedly lower in d1 plants under low-light conditions compared with control conditions; the opposite pattern was observed for INV activity in OE-1 plants.

The activities of SUS and INV are regulated by several genes, and the expression levels of these genes were determined. The expression of *SUS1* was up-regulated in WT and OE-1 plants by low-light stress, and its expression was higher in OE-1 plants than in WT plants (Fig. 4B, a). No significant difference in *SUS1* expression was observed in d1 plants between control and low-light conditions. Low-light stress did not affect the relative expression levels of *SUS3* in WT and d1 plants; however, *SUS3* expression was significantly increased in OE-1 plants by low-light stress (Fig. 4B, b). The expression of *INV1* and cell-wall invertase 1 (*CIN1*) was highest in OE-1 plants, followed by d1 plants and WT plants, under low-light conditions (Fig. 4B, c and e). No differences in the relative expression levels of *INV2* and *CIN2* were observed in WT plants between control and low-light conditions (Fig. 4B, e and f). The relative expression levels of *INV2* and *CIN2* were substantially decreased by low-light stress in d1 plants; however, low-light stress increased the expression of these two genes in OE-1 plants.

### *RGA1* improves energy status in the pistil under low-light conditions

The mitochondrial respiratory electron transport chain complexes were studied to clarify the ability of *RGA1* to contribute to energy production in the pistil under low-light conditions. There were no differences in the activities of complex I, IV, and V in WT plants between control and low-light conditions; however, marked decreases and increases in the activities of these complexes were observed in d1 and OE-1 plants, respectively, under low-light conditions compared with control conditions (Fig. 5, a, b and c).

**Figure 5.**
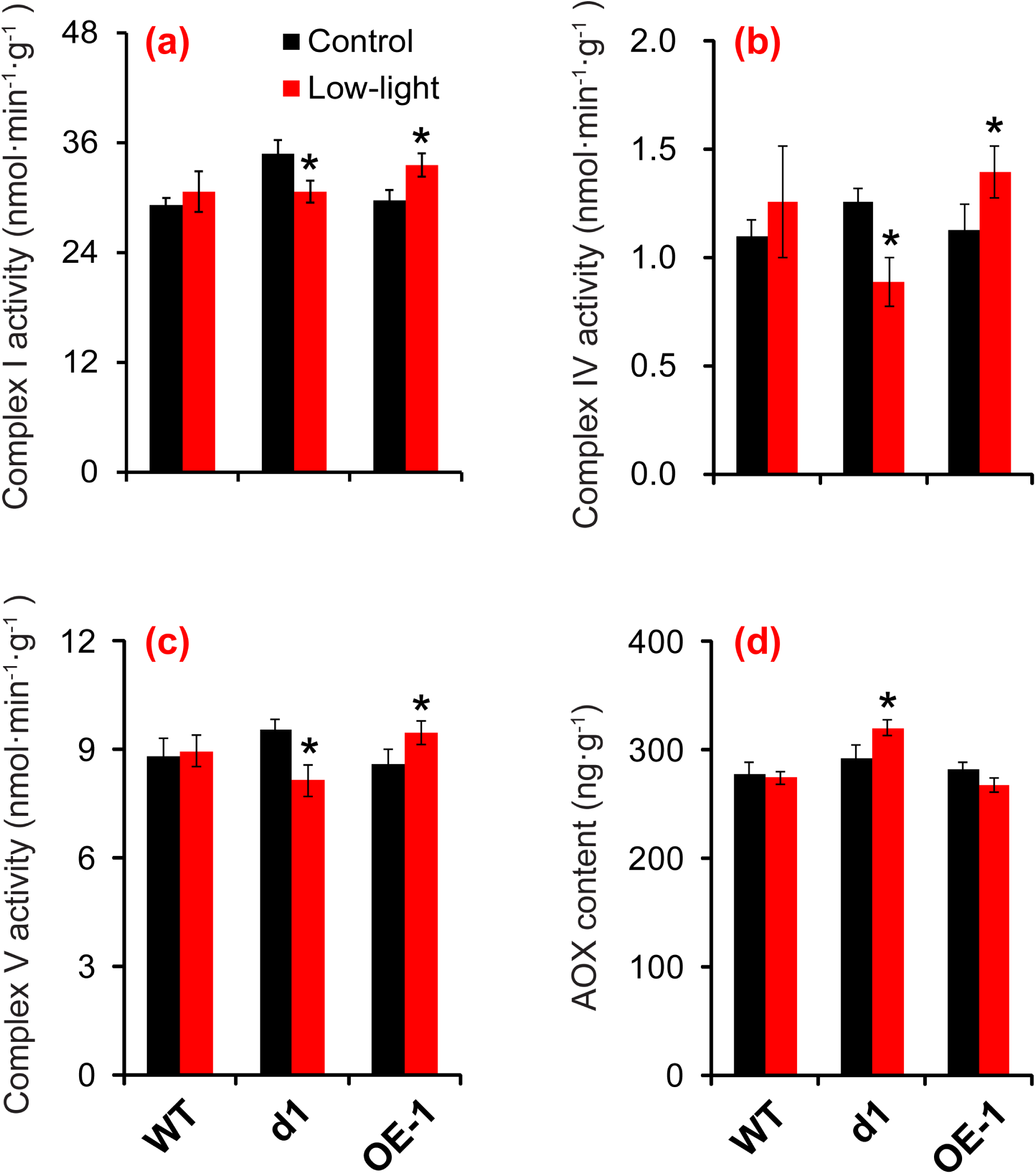
Effects of low-light stress on mitochondrial respiratory electron transport in the pistil of rice at anthesis. **a**, Complex I activity; **b**, Complex IV activity; **c**, Complex V activity; **d**, AOX content. AOX, Alternative oxidase. The mean values and standard errors in the figures correspond to data from three replicates. Student’s *t*-tests were conducted to evaluate the significance of differences between treatments within single genotypes. * indicates P-value < 0.05.

The content of AOX and PARP is an important factor affecting the energy production efficiency of plants. No difference in AOX content was observed in WT plants between control and low-light conditions, and AOX content was higher in d1 plants under low-light conditions compared with control conditions (Fig. 5, d). By contrast, the AOX content in OE-1 plants was lower under low-light conditions compared with control conditions. Low-light stress had little effect on the PARP content in WT plants (Fig. S5). However, the PARP content of d1 plants was higher under low-light conditions than under control conditions, and the content of PARP decreased under low-light conditions compared with control conditions in OE-1 plants.

### *RGA1* optimizes energy allocation in the pistil under low-light stress

The activity of SOD, POD, and CAT was significantly higher in d1 plants under low-light conditions compared with control conditions, and the opposite pattern was observed in OE-1 plants (Fig. 6A, a, b and c). There was no noticeable difference in the activity of these enzymes in WT plants between control and low-light conditions (Fig. 6A, a, b, c, and d). There was no marked difference in the activity of APX in WT and d1 plants between control and low-light conditions; APX activity was significantly lower in OE-1 plants under low-light conditions compared with control conditions (Fig. 6A, d).

**Figure 6.**
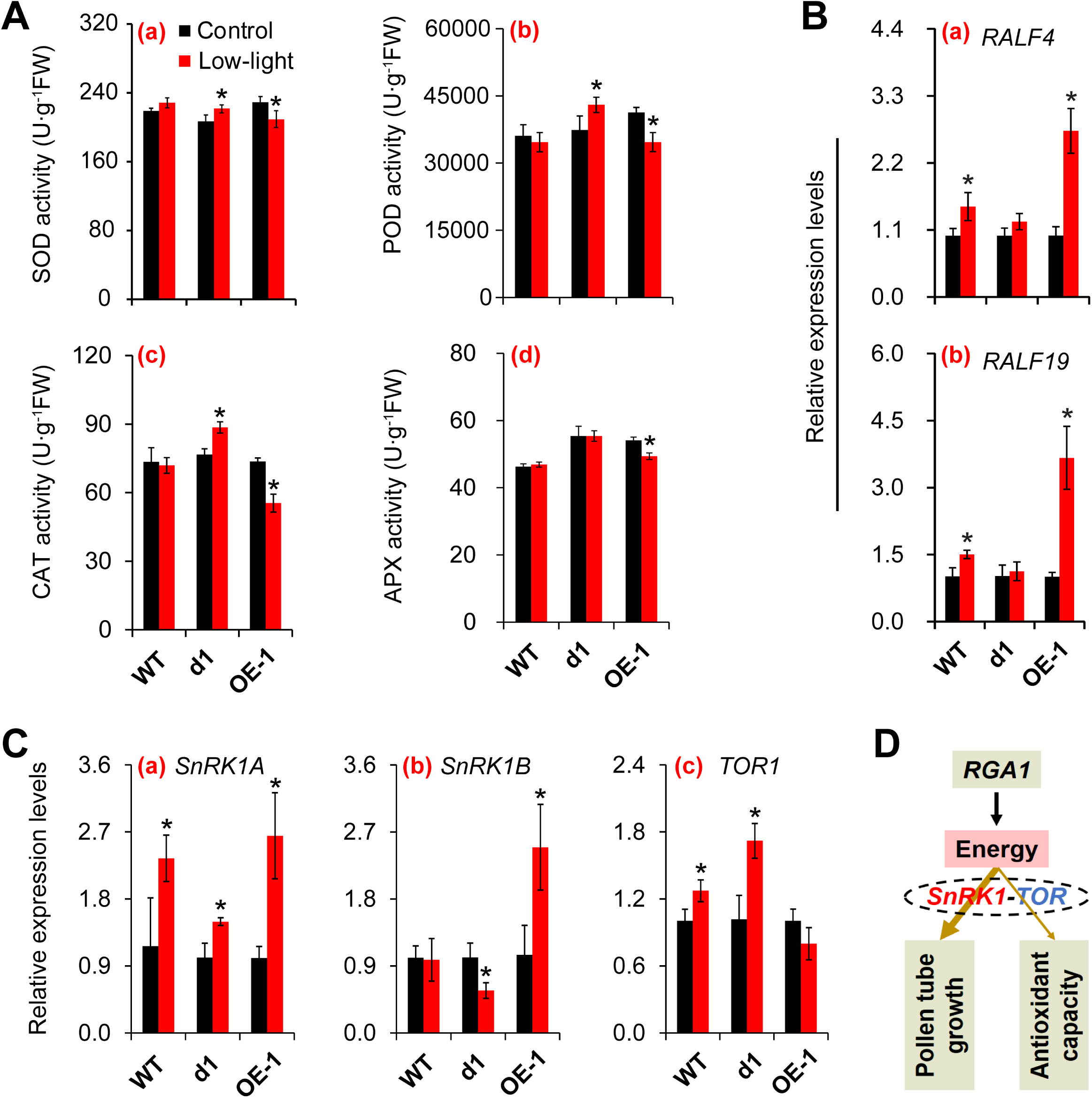
Effect of low-light stress on energy allocation in the pistil of rice at anthesis. **a**, SOD activity; **b**, POD activity; **c**, CAT activity; **d**, APX activity. **B**, Changes in *RALFs* genes in pistil under low-light condition. **C**, Changes in expressions of *SnRK1* and *TOR* genes in pistil of rice under low-light condition. **a**, *SnRK1A*; **b**, *SnRK1B*; **c**, *TOR1*. **D**, Schematic diagram of *RGA1* regulating energy allocation through *SnRK1* and *TOR*. SOD, Superoxide dismutase; POD, Peroxidase; CAT, Catalase; APX, Ascorbate peroxidase. The mean values and standard errors in the figures correspond to data from three replicates. Student’s *t*-tests were conducted to evaluate the significance of differences between treatments within single genotypes. * indicates P-value < 0.05.

The process of pollen tube elongation is controlled by RALF family peptides; thus, the relative expression levels of *RALF4* and *RAL19* were determined. There was no difference in the relative expression level of *RALF4* between control and low-light conditions in d1 plants. However, there was a notable increase in the expression of *RALF4* in WT and OE-1 plants under low-light conditions, and the increase in *RALF4* expression under low-light conditions relative to control conditions was more pronounced in WT plants than in OE-1 plants (Fig. 6B, a). *RALF19* expression was activated by low-light stress, and it was most highly up-regulated in OE-1 plants, followed by WT and d1 plants (Fig. 6B, b).

SnRk1 and target of rapamycin (TOR) are energy sensors that play key roles in regulating energy metabolism and allocation in plants under various environmental conditions. The expression of *SnRK1A* was activated by low-light stress, and the highest expression of *SnRK1A* was observed in OE-1 plants, followed by WT and d1 plants (Fig. 6C, a and b). The relative expression levels of *SnRK1B* in WT plants were not affected by low-light conditions (Fig. 6C, b). However, low-light stress significantly decreased the relative expression level of *SnRk1B* in d1 plants. In OE-1 plants, the expression of this gene was significantly up-regulated by low-light stress. The expression patterns of *TOR1* were opposite those of *SnRk1A* and *SnRK1B*. Under low-light conditions, the relative expression levels of *TOR1* were highest in d1 plants, followed by WT plants and OE-1 plants (Fig. 6C, c). These findings indicate that although energy is typically mainly allocated to the process of pollen tube elongation in the pistil of OE-1 plants (Fig. 6D), more energy was allocated to enhancing antioxidant capacity rather than pollen tube elongation in the pistil of d1 plants under low-light conditions.

### Effect of sucrose, Na_2_SO_3_, and Na_2_VO_4_ on spikelet fertility and energy metabolism under low-light conditions

According to the above findings, INV and ATPase might play important roles in maintaining energy homeostasis in rice plants under low-light conditions. To confirm this hypothesis, sucrose, Na_2_SO_3_, and Na_2_VO_4_ were sprayed on rice plants, and the spikelet fertility, the content of ATP and ATPase, and the activities of SUS and INV were determined. Low-light stress induced spikelet sterility in these three genotypes (Fig. 7). These reductions in spikelet fertility were partly reversed when the plants were treated with sucrose and Na_2_SO_3_, and marked differences were observed in WT and d1 plants. By contrast, Na_2_VO_4_ markedly reduced the spikelet fertility of WT and OE-1 plants under low-light conditions but had little effect on d1 plants.

**Figure 7.**
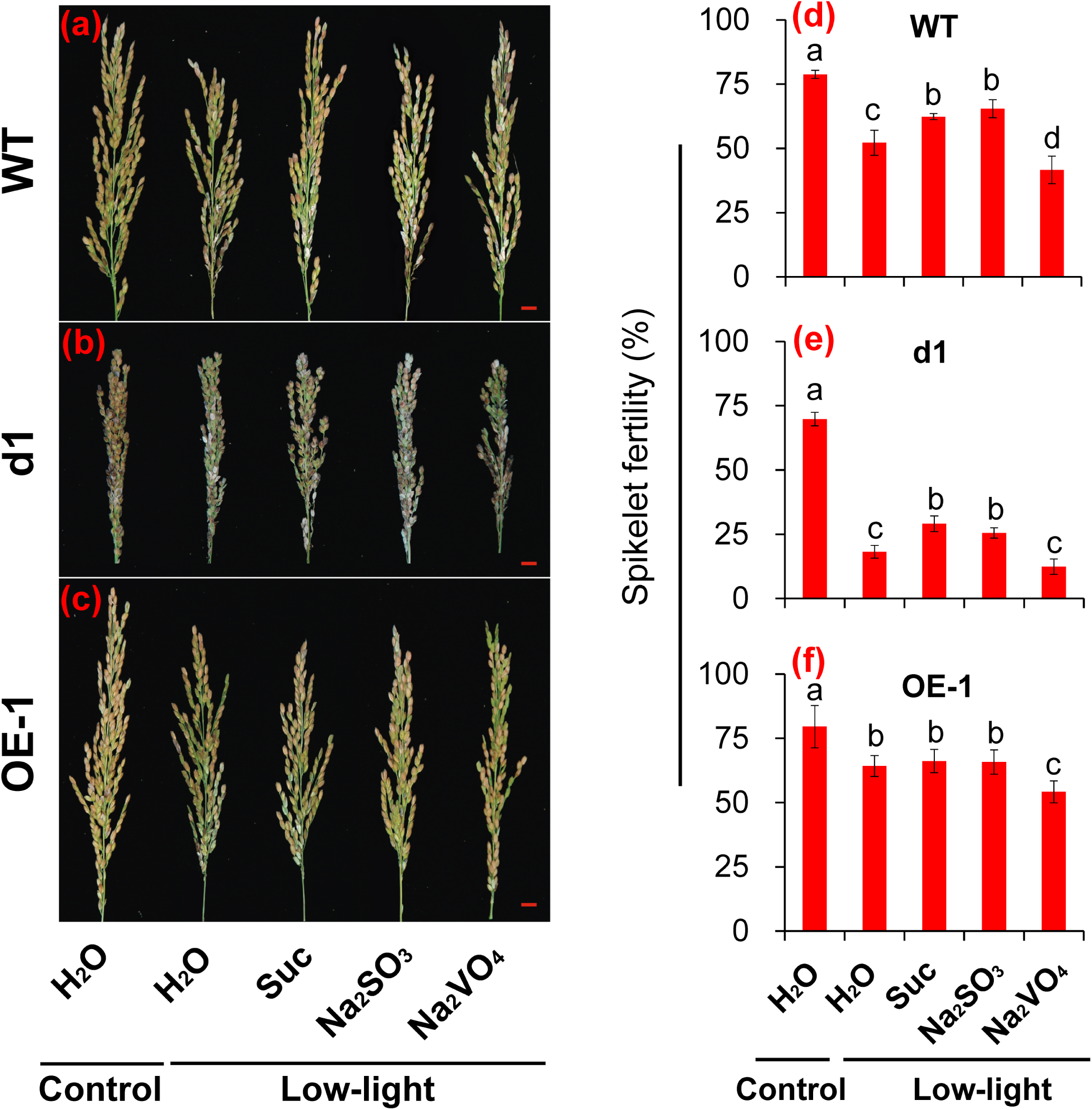
Effect of sucrose, Na_2_SO_3_, and Na_2_VO_4_ on the spikelet fertility of rice under low-light conditions. **a-c**, The panicle morphology of rice. Bar = 1 cm; **d-f**, Spikelet fertility. The mean values and standard errors in the figures correspond to data from five replicates. One-way analysis of variance (ANOVA) was conducted to compare the difference with a least significant difference test (LSD) at P ≤ 0.05 for the data of spikelet fertility within one genotype. Different letters indicate significant differences among different treatments within one genotype.

Changes in ATP, ATPase, INV, and SUS varied among plants treated with H_2_O, sucrose, Na_2_SO_3_, and Na_2_VO_4_ under low-light conditions (Fig. 8). There was no marked difference in the content of ATP and ATPase among H_2_O, sucrose, and Na_2_SO_3_ treatments in WT and OE-1 plants (Fig. 8, a, c, d, and f); large increases in the content of ATP and ATPase were observed in d1 plants under sucrose and Na_2_SO_3_ treatment compared with H_2_O treatment (Fig. 8, b and e). By contrast, Na_2_VO_4_ had little effect on the content of ATP and ATPase in d1 plants, but Na_2_VO_4_ greatly reduced the content of ATP and ATPase in WT and OE-1 plants. Under low-light conditions, no differences in INV and SUS activity were observed among plants treated with H_2_O, sucrose, and Na_2_SO_3_ in WT and OE-1 plants; however, large decreases in INV and SUS activity were observed in plants treated with Na_2_VO_4_ (Fig. 8, g–l). INV and SUS activity was higher under sucrose treatment than under H_2_O treatment, and no significant differences in INV and SUS activity were observed between Na_2_SO_3_ treatment and H_2_O treatment (Fig. 8, h and k).

**Figure 8.**
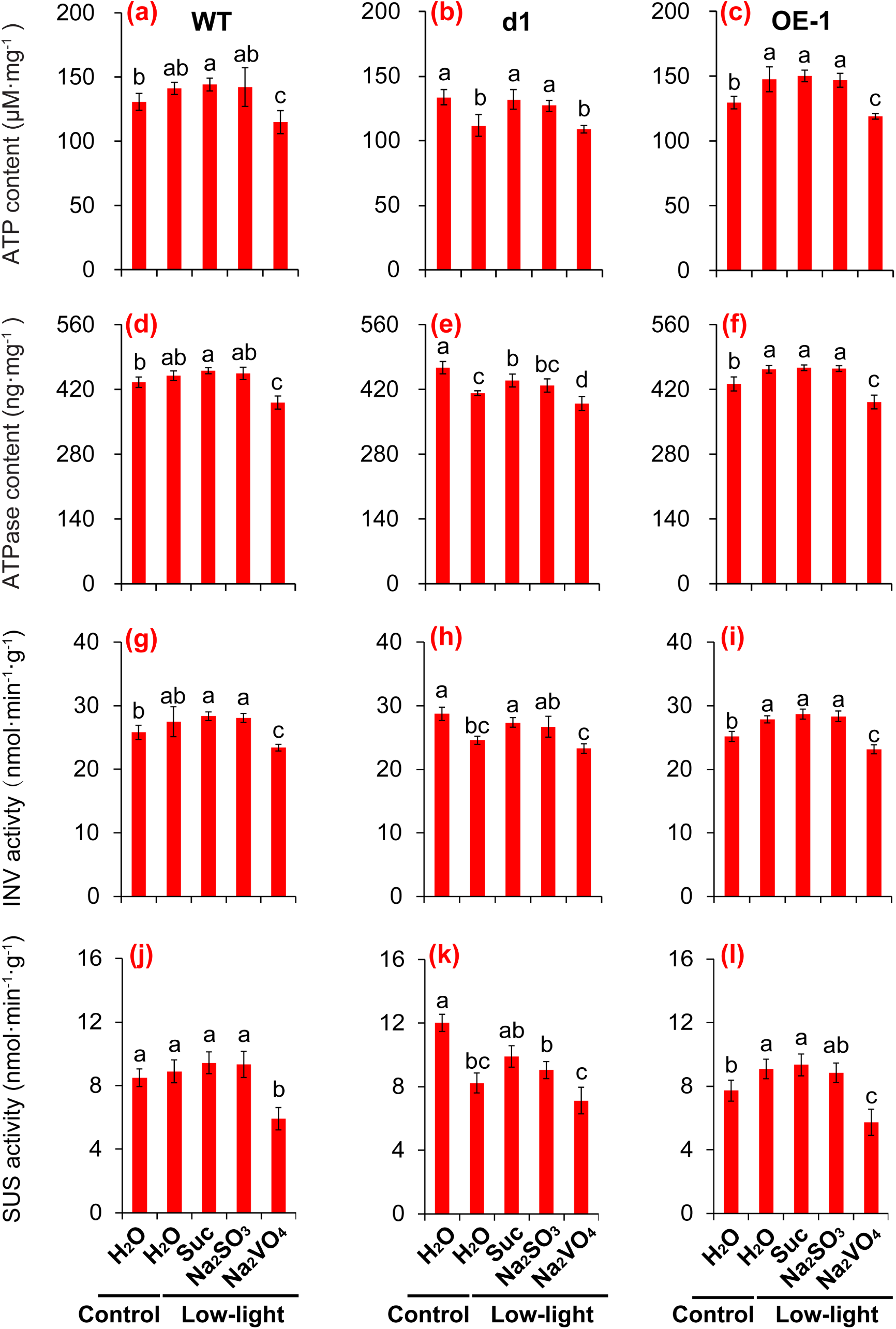
Effect of sucrose, Na_2_SO_3_, and Na_2_VO_4_ on the energy status and sucrose metabolism in the pistil of rice under low-light conditions. The mean values and standard errors in the figures correspond to data from three replicates. One-way analysis of variance (ANOVA) was conducted to compare the difference with a least significant difference test (LSD) at P ≤ 0.05 for the data of spikelet fertility within one genotype. Different letters indicate significant differences among different treatments within one genotype.

## Discussion

### *RGA1* confers low-light tolerance in rice plants

The gene *RGA1* was first identified in rice in 1995 (Ishikawa et al., 1995). This gene is not only involved in regulating plant growth and development but also in the responses to various types of abiotic stress, such as drought (Ferrero-Serrano & Assmann, 2016), cold (Ma et al., 2015), salt (Jangam et al., 2016), high-light (Ferrero-Serrano et al., 2018), and heat stress (Kan et al., 2022). Spikelet sterility was higher in *RGA1* mutant plants under low-light conditions at anthesis compared with WT and OE-1 plants (Fig. 1), suggesting that *RGA1* enhances low-light tolerance in rice plants; this is a novel finding that has not been documented in previous studies. *RGA1* mutant plants had a greater capacity to dissipate excess irradiance, which could enhance photoavoidance and photoprotection and thus reduce sustained photoinhibitory damage (Ferrero-Serrano et al., 2018). These findings suggest that the functions of *RGA1* in rice plants differ between low-light and high-light stress. Low-light stress significantly decreases the net photosynthetic rate in plants (Slattery et al., 2018; Kono et al., 2020), and less pronounced decreases in the net photosynthetic rate were observed in *RGA1* mutant plants compared with WT and OE-1 plants (Fig. S2). Thus, *RGA1* might confer low-light tolerance in rice plants by improving sugar and energy metabolism rather than light-use efficiency or photosynthetic assimilate production.

Recently, reports on regulators of G protein signaling (RGS1) combined with G protein complex to sense apoplast glucose concentration for signal transduction have gradually emerged (Peng et al., 2018; Wang et al., 2022). A lack of light reduces the output of photosynthesis in the leaves of rice plants, and the metabolism of sugar decreases sugar levels in plants. Wang et al. (2022) showed that the RGS1/G protein complex senses changes in glucose concentrations in low light and regulates tomato disease resistance by signaling, and Gα negatively regulates light-mediated defense. In our study, *RGA1* positively regulated low-light tolerance in rice, suggesting that its responses to biotic and abiotic stress differ. Nonetheless, it was determined that RGS1 senses sugar levels inhibited by low-light stress through binding to RGA1 (Gα subunit) to transmit signals to downstream effectors for defense responses. Our study was focused on elucidating the downstream physiological defense mechanisms; however, the decoding of these signals in this process still requires additional study.

### *RGA1* improves energy status to promote pollen tube elongation under low-light conditions

Low-light stress can induce spikelet sterility because of the poor pollen viability and low anther dehiscence (Chen et al., 2019; Deng et al., 2021). However, no differences in pollen viability, anther dehiscence, number of pollen grains on the stigma, and pollen germination were observed between control and low-light conditions in all three rice genotypes (Fig. 2A; Fig. S3 and S4). The ability of the pollen tube to reach the ovule was the main factor contributing to the higher spikelet fertility of WT and OE-1 plants compared with d1 plants under low-light conditions (Fig. 2A, i). These inconsistent findings might be related to the magnitude or duration of stress. Plants were only exposed to low-light stress for 7 d at anthesis, and the status of the pollen tubes was determined at 4–5 d after stress; by contrast, plants were exposed to shading stress for 45 d in the study of Deng et al. (2021). The heterotrimeric G protein is involved in pollen germination and pollen tube elongation (Zhang et al., 2005; Wu et al., 2007). Suppression of the heterotrimeric G protein beta-subunit results in an aberrant anther shape and the production of non-germinating pollen in tobacco (Peskan-Berghöfer et al., 2005). However, the rate of pollen germination and pollen tuber growth into ovule of all the three genotypes exceeded 40% under natural conditions, though lower values were showed in d1 plants than those of WT and OE-1 plants (Fig. 2A, h and i). This means that *RGA1* functioned in pollen germination and pollen tube elongation under low-light conditions rather than under natural conditions, and this is inconsistent with the results of previous studies (Zhang et al., 2005; Peskan-Berghöfer et al., 2005; Wu et al., 2007).

During the pollination and fertilization process in plants, cell–cell communication is required for pollen tube elongation, as this provides the directional guidance needed for pollen tubes to reach the ovule (Dresselhaus, et al., 2016; Zhang et al., 2017). Signaling molecules such as H_2_O_2_ and Ca^2+^, as well as energy, play important roles in this process (Zhang et al., 2018a; Kim et al., 2021; Zhou et al., 2021). G proteins modulate Ca^2+^ and H_2_O_2_ to regulate pollen germination, pollen tube elongation, and stomatal aperture (Zhang et al., 2005; Peskan-Berghöfer et al., 2005; Zhang et al., 2011). We found that H_2_O_2_ levels were higher in the pistils of d1 plants than in WT and OE-1 plants under low-light conditions (Fig. 2C, a). However, high H_2_O_2_ levels in the pistil might act as a signaling molecule rather than as a toxic substance. This hypothesis was confirmed by the notable reduction in the content of MDA in all three genotypes under low-light stress (Fig. 2C, b); the MDA content is generally considered an indicator of cellular damage (Islam et al., 2019; Yu et al., 2020). Moreover, low-light stress had no effect on the gradient trend of the fluorescence intensity of ROS in the pistil, and it gradually increased from the stigma to the style and the ovary (Fig. 2B). This suggests that the disturbance of pollen tube elongation by low-light stress was not the main factor causing the spikelet sterility of d1 plants. The content of ATP and ATPase was lower in d1 plants than in WT and OE-1 plants under low-light stress (Fig. 2D, a and b). Energy status is important for pollen tube elongation in the pistil, especially under abiotic stress, as pollen tube elongation is an energy-demanding process (Cárdenas et al., 2006; Certal et al., 2008). Thus, energy deficits caused by low-light stress might be the main cause of the cessation of pollen tube growth in the pistils of d1 plants (Jiang et al., 2020; Hoffmann et al., 2020).

### *RGA1* optimizes sucrose transport and metabolism in rice plants under low-light stress

A sufficient supply of carbon to the spikelets is key for ensuring the reproductive growth of rice, especially under adverse conditions (Zhang et al., 2018b; Islam et al., 2019; Chen et al., 2022a). Shading or low-light stress inhibits the transfer of carbon assimilates to the sink organs of crops, such as rice, maize, and wheat, and thus yield (Kobata et al., 2000; Chen et al., 2020; Shao et al., 2021). We found that *RGA1* could alleviate the impairment of sucrose transport and metabolism by low-light stress to improve the source-sink relationship in the pistil, and SUTs (Hackel et al., 2006), SWEETs (Sosso et al., 2015; Yang et al., 2018), and INV (Ruan, 2014; Jiang et al., 2020) play key roles in this process (Fig. 3 and 4). Sugar signaling and phytohormones regulate the distribution of carbon assimilates in crops, especially under abiotic stress (Yu et al., 2015; Zhang et al., 2018b; Chen et al., 2022b). The heterotrimeric G protein has been found to be involved in sugar and phytohormone signaling in plants (Peng et al., 2018; Zhang et al., 2021). This might suggest that *RGA1* can improve the source–sink relationship by regulating sugar and phytohormone signaling under low-light conditions.

Sucrose can be hydrolyzed to glucose and fructose by SUS and INV (Wang et al., 1993; Roitsch & Gonzalez, 2004). Shading and low-light stress can reduce SUS and INV activity in the potato tuber cambium (Chen & Setter, 2003). In d1 plants, no difference in SUS activity was observed between control and low-light conditions, and a notable decrease in INV activity was observed under low-light stress (Fig. 4A, e and f). Similar results have been observed in rice plants under heat stress at anthesis; specifically, INV, rather than SUS, maintains energy homeostasis in rice spikelets to enhance pollen tube elongation in the pistil (Jiang et al., 2020). This suggests that SUS and INV have different functions in plants, especially under abiotic stress (Wang et al., 1993; Barratt et al., 2009; Stein & Granot, 2019); this has been discussed in detail in Jiang et al. (2020). Recent studies have shown that SUS is not involved in starch synthesis in Arabidopsis leaves (Fünfgeld et al., 2022). However, SUS and INV activity, and the relative expression levels of related genes, were significantly increased in OE-1 plants under low-light stress (Fig. 4A, e and f; Fig. 4B), suggesting that more research is needed to characterize the mechanisms underlying these patterns.

Sucrose is converted into glucose and fructose by SUS and INV, which then enters glycolysis, followed by the tricarboxylic acid cycle, the pentose phosphorylation pathway, and finally oxidative phosphorylation in mitochondria to generate ATP (Plaxton & Podestá, 2006). In this study, *RGA1* was shown to enhance the activities of complex I, IV, and V as well as decrease the content of AOX and PARP (Fig. 5 and Fig. S5), which directly reflects ATP production efficiency in the mitochondria (Plaxton & Podestá, 2006; De Block & Van Lijsebettens, 2011; O’Leary et al., 2020). G protein-coupled receptors can sense metabolic activity or levels of energy substrates to control the secretion of metabolic hormones or regulate the metabolic activity of particular cells in animals. Furthermore, the heterotrimeric G protein is a component of a signal transduction pathway for ATP release from erythrocytes (Olearczyk et al., 2004a, b). In plants, the heterotrimeric G protein is involved in ATP-promoted stomatal opening (Hao et al., 2012). This suggests that the heterotrimeric G protein might regulate the activities of these complexes to increase the energy production efficiency of plants under low-light conditions.

### *RGA1* optimizes energy allocation in the pistil under low-light stress

Life is a process of energy consumption, and protein turnover, metabolic activity, ion transport, futile cycling, sucrose transport, the uptake and utilization of nitrogen, and the response to abiotic stress require large amounts of energy in plants (Amthor et al., 2019). Thus, optimizing energy allocation and maintaining energy homeostasis are major challenges for all living organisms under abiotic stress (De Block & Van Lijsebettens, 2011; Dröge-Laser & Weiste, 2018). The active transport of assimilates, increases in antioxidant capacity, and the accumulation of heat shock proteins are energy-demanding processes that play key roles in abiotic adversity defense (Yu et al., 2020). The accumulation of heat shock proteins was significantly decreased by low-light stress (Fig. S6), but the antioxidant capacity, including SOD, POD, and CAT activities, was significantly increased in d1 plants under low-light stress (Fig. 6A). This suggests that the energy in the pistil of d1 plants is mainly allocated to antioxidant capacity; consequently, the energy allocated to the process of pollen tube elongation is limited (Rottmann et al., 2016; Goetz et al., 2017; Ge et al, 2019), and this results in the cessation of pollen tube growth and spikelet sterility (Zhang et al., 2018a). To test this hypothesis, we examined the *RALF4* and *RALF19* genes, which are mainly responsible for pollen tube elongation in the pistil (Ge et al., 2017; Li & Yang, 2018). The relative expression levels of *RALF4* and *RALF19* were significantly induced by low-light stress in WT and OE-1 plants, but no differences in the expression levels of these genes were observed in d1 plants between control and low-light conditions (Fig. 6B). This suggests that *RGA1* can improve the allocation of energy in the pistil to enhance pollen tube elongation under low-light conditions.

Respiration is divided into growth respiration and maintenance respiration (Amthor et al., 2019). In growth respiration, energy mainly contributes to plant biomass accumulation, whereas energy is used to maintain cellular homeostasis to ensure cell survival in maintenance respiration (Amthor et al., 2019; Li et al., 2021). The balance between these two processes determines yield and quality under abiotic stress by regulating energy utilization efficiency (Lee & Millar, 2016). Two protein kinase complexes, SnRK1 and TOR, are central hubs in metabolic regulation and have antagonistic effects on plant growth (Baena-González & Hanson, 2017; Margalha et al., 2019). SnRk1 and TOR also function in energy allocation in the pistil (Li et al., 2017). Under low-light conditions, patterns of SnRK1 and TOR activity in these genotypes varied (Fig. 6C), and this is consistent with the results of previous studies (Baena-González & Hanson, 2017). This suggests that *RGA1* can achieve a balance between growth and maintenance respiration by regulating the activities of SnRK1 and TOR in the pistil under low-light conditions, which can enhance energy utilization efficiency and provide sufficient energy for the process of pollen tube elongation into the ovule. However, the mechanism by which SnRK1 and TOR are regulated by *RGA1* in plants remains unclear.

In addition to the direct sensing of glucose signals by the RGS1/G protein complex, SnRK1 and TOR can indirectly respond to sugar signals by sensing energy and nutritional status; SnRK1/TOR signaling is thus a key sugar signaling pathway (Chen et al., 2022b). Our results suggest that SnRK1/TOR signaling and RGS1/G protein signaling appear to be interconnected. Hexokinase (HXK) is also considered an important sugar signaling sensor (Kim et al., 2013). In this experiment, a large increase in the expression of *HXK10-1* was observed in WT and OE-1 plants under low-light stress, and this was consistent with patterns of SnRK/TOR signaling under low-light stress (Fig. S7, b). In a previous study, crosstalk between G protein signaling and hexokinase was shown to mediate glucose signaling in plants (Huang et al., 2015). Whether there is convergence in the direct and indirect sugar signaling pathways remains unclear (Chen et al., 2022b). Although our data suggest that the RGS1/G protein signaling, SnRK1/TOR signaling, and HXK signaling appear to be related, the available data do not provide strong support for this hypothesis.

## Conclusion

Low-light stress significantly decreased the spikelet fertility of rice at anthesis. *RGA1* partly alleviated low light-induced spikelet sterility by preventing the cessation of pollen tube growth in the pistil through its effects on the distribution of carbohydrates, energy metabolism, and energy allocation (Fig. 9). The ROS distribution in pistils of the three genotypes was stable, and the energy status of WT and OE-1 plants was improved compared with that of d1 plants under low-light conditions, suggesting that *RGA1* enhanced pollen tube elongation in the pistil by maintaining energy homeostasis, rather than by regulating signaling pathways involved in guiding the growth of the pollen tube. No differences in assimilate production were observed among these three genotypes under low-light conditions, but greater amounts of assimilates were transported to the panicle in WT and OE-1 plants than in d1 plants. In this process, greater increases in the expression levels of *SUTs* and *SWEETs*, as well as SUS and INV activities, were observed in WT and OE-1 plants compared with d1 plants. This indicated that *RGA1* could enhance sucrose transport and metabolism in the pistil under low-light conditions. *RGA1* optimizes the efficiency of energy production by increasing the activities of mitochondrial complex I, IV, and V and reducing the AOX content. *RGA1* also improved energy allocation by regulating the expression levels of the *SnRK1* and *TOR* genes, which enhanced energy utilization efficiency and provided sufficient energy for pollen tube elongation in the pistil. In sum, *RGA1* allowed sufficient energy to be available for pollen tube elongation in the pistil under low-light conditions by promoting sucrose unloading in the pistil as well as improving energy production, allocation, and utilization.

**Figure 9.**
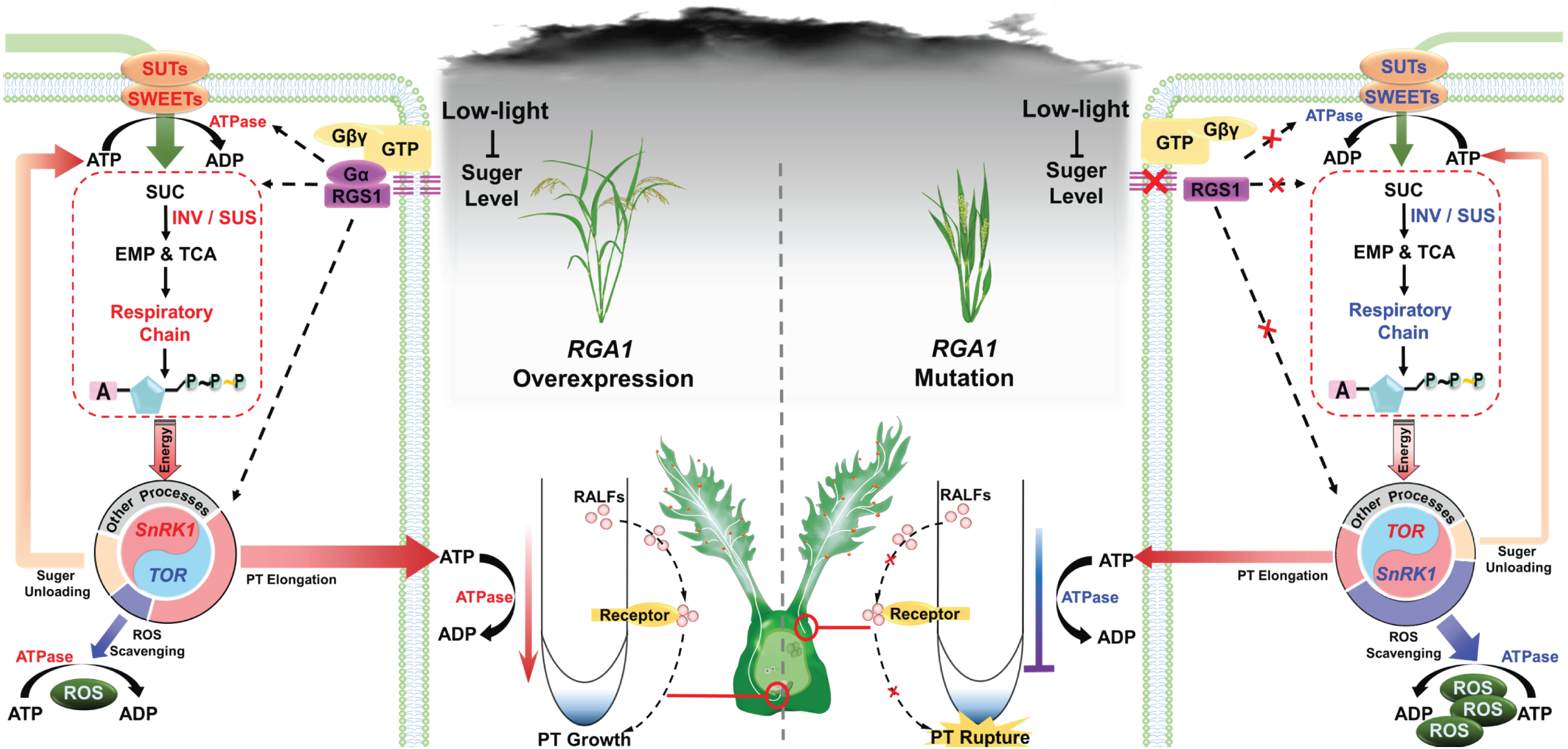
Descriptive model of the function of *RGA1* in pollen tube elongation in the pistil under low-light stress. **Left:** G protein signaling 1 (*RGS1*) senses the inhibition of sugar and energy levels by low-light stress and interacts with the Gα subunit (*RGA1*) to activate the G protein-mediated low-light tolerance response. *RGA1* activates SUTs, SWEETs, INV, SUS, mitochondrial respiratory electron transport chain complexes, and ATPase under low-light conditions. This can enhance sucrose transport and metabolism in the pistil and thus energy status. Furthermore, *RGA1* activates *SnRK1* antagonism and represses *TOR* to improve energy allocation, most of which is used for active sugar transport and pollen tube elongation. Sufficient energy ensures pollen tube elongation and sustains the secretion of RALFs to maintain pollen tube integrity until the ovary is reached. **Right:** Deficiencies of *RGA1* impede the G protein-mediated low-light tolerance response. In d1 plants, the expression of *SUTs* and *SWEETs* and the activities of INV, SUS, mitochondrial respiratory electron transport chain complexes, and ATPase are repressed by low-light stress, which negatively affects energy status. *TOR* is activated, and *SnRK1* is repressed, which increases the allocation of energy to enhance the antioxidant capacity to reduce excess ROS rather than pollen tube elongation. A lack of available energy inhibits pollen tube elongation and can even cause the rupture of the pollen tube. In the figure, red font indicates activation, blue font indicates inhibition, and dashed arrows indicate possible direct mediation.

## Materials and methods

### Plant growth conditions and treatments

Zhonghua 11 (wild type, WT), the Gα heterotrimeric G protein subunit mutant (*RGA1* mutant, d1), and transgenic rice lines overexpressing *RGA1* (OE-1) were sown in buckets (10-cm radius; 20-cm height) at the China National Rice Research Institute (CNRRI), Hangzhou, China. Rice seeds were soaked for 48 hours at 30°C and incubated for 24 hours at 35°C to germination. After the rice seedlings had grown four leaves, three of the plants were transplanted into each pot and filled with paddy soil until flowering. Thereafter, rice plants were divided into two groups in a plant growth chamber. An automatic temperature control system was used to control both temperature and relative humidity. One group of rice plants was subjected to a low-light intensity of 30 μmol·m^-2^·s^-1^ for 7 d at anthesis, and the control group was subjected to a light intensity of 300 μmol·m^-2^·s^-1^. Both groups were maintained at 70– 80% relative humidity and normal temperature conditions (28°C/23°C: day/night). The flowering spikelets, including the pistil, were sampled 3–4 d after exposure to low-light stress.

According to the above findings, sucrose, sodium sulfite (Na_2_SO_3_), and sodium ortho-vanadate (Na_2_VO_4_) were sprayed on rice plants under low-light conditions. Sucrose was used as the INV activator (Jiang et al., 2020), and Na_2_SO_3_ and Na_2_VO_4_ were used as activators and inhibitors of ATPase, respectively (Oliveira et al., 2004). These chemicals (0.1% sucrose, 5 mM Na_2_SO_3_, or 5 mM Na_2_VO_4_) were sprayed on rice plants 2 d after exposure to low-light stress. The spikelets, including pistils, collected 5 d after low-light stress were used to determine the content of ATP and ATPase as well as the activities of SUS and INV.

### Measurement of grain yield, yield components, dry matter weight, and net photosynthesis

At maturity, rice plants were harvested from low-light stress and control groups to determine the grain yield per plant, spikelet fertility, kernel weight, and dry matter weight. The spikelet fertility was calculated as FG/(FG+AG)×100% (FG, Filled grain number; AG, Aborted grain number). Approximately 100 grains were spread on the grass plate of a flatbed scanner and scanned. The kernel weight was determined using SC-G software (Hangzhou Wanshen Detection Technology Co., Ltd., Hangzhou, China). The rice plants were divided into leaf, sheath-stem, and panicle and then were dried at 85 °C for 48 h to determine the weight and distribution of dry matter.

Net photosynthesis was determined using a Li-COR 6400 portable photosynthesis system (Li-COR Biosciences Inc., Lincoln, NE, USA) under the following conditions: photosynthetic photon flux density of 1,200 μmol·m^−2^·s^−1^; ambient CO_2_ (450 μmol·mol^−1^); 6 cm^2^ leaf area; 500 μmol·s^−1^ flow speed; and temperatures according to the treatment.

### Measurements of pollen viability, anther dehiscence, pollen germination, and pollen tube elongation in the pistil

Pollen viability was determined following the method of Islam et al. (2019). Non-flowering spikelets on the fourth to sixth primary branches of panicle-containing anthers were collected on Day 3 of the low-light stress treatment. More than 50 flowering spikelets were collected on days 3 to 4 of the low-light stress treatment, and anther dehiscence was observed using a stereomicroscope (SteREO Lumar.V12, Zeiss, Germany). More than 50 flowering spikelets were collected on days 3 to 4 of the low-light stress treatment, and the tissues were subsequently stained following the method of Zhang et al. (2018a). The number of pollen grains in the pistil was determined, and pollen germination and pollen tube elongation were observed using fluorescence microscopy (DM4000B, Leica, Wetzlar, Germany) at 350 nm.

### Measurement of the levels of reactive oxygen species (ROS) in the pistil

Pollinated pistils were collected from flowering spikelets and immediately immersed in a reaction solution of 2 μM DCFH-DA as described in Fu et al. (2016). After 30 min of incubation at room temperature, the pistils were washed with 0.01 M PBS (pH 7.0); images were acquired and the fluorescence intensity was measured using a fluorescence microscope (DM4000B, Leica, Wetzlar, Germany).

### Measurements of lipid peroxidation (malondialdehyde, MDA) and H_2_O_2_ content

Approximately 0.01 g of frozen pistil was homogenized in 2 mL of 5% trichloroacetic acid, and then the MDA content was estimated by measuring the concentration of thiobarbituric acid-reactive substances (Dhindsa et al. 1981). The H_2_O_2_ content was determined following the method of Brennan and Frenkel (1977) with some modifications; approximately 0.01 g of frozen pistil was homogenized in 4 ml of 10 mM 3-amino-1,2,4-triazole and then centrifuged for 25 min at 6,000 g. Thereafter, 0.1% titanium tetrachloride in 1 ml of 20% H_2_SO_4_ was added to 2 ml of the supernatant. The reaction solution was further centrifuged to remove undissolved materials, and the absorbance was recorded at 410 nm.

### Carbohydrate content measurements

The total soluble sugar and starch content was determined using the sulfuric acid anthrone colorimetric method with modifications (Dubois et al., 1956). Sucrose, glucose, and fructose were extracted using the same method that was used for the extraction of total soluble sugars, and the content of sucrose, glucose, and fructose was determined using the method of Zhang (1977) with minor modifications (Zhang et al., 2018b). The content of total non-structural carbohydrates (NSC) was calculated as the sum of the content of soluble sugars and starch.

### Measurement of SUS and INV activities

Frozen pistils (0.01 g) were ground to a fine powder in liquid nitrogen and then homogenized in extraction buffer (20%) containing 50 mM HEPES (pH 7.5), 5 mM MgCl_2_, 1 mM ethylene diamine tetra acetic acid disodium salt (EDTA-Na_2_), 0.5 mM dithiothreitol (DTT), 1% Triton X-100, 2% polyvinyl pyrrolidone, and 10% glycerol. The supernatant was immediately desalted on a Sephadex G-25 column equilibrated with extraction buffer at 4°C. The filtrate was then used to determine the SUS and INV activities with a test kit (Comin Biotechnology Co., Ltd., Suzhou, China).

### Measurements of mitochondrial complex activities

Frozen pistils (0.01 g) were ground, and mitochondria were extracted in an ice bath with reagents in the mitochondrial complex activity assay kits; the extract was subjected to 30 cycles of ultrasonic treatment (power, 20%; ultrasonic time, 3 s; and interval time, 10 s). Mitochondrial complex activity assay kits were used to determine the activity of the mitochondrial complexes following the manufacturer’s instructions (Comin Biotechnology Co., Ltd., Suzhou, China).

### Measurements of the content of ATP, ATPase, and AOX and PARP

Frozen pistils (0.01 g) were ground to a fine powder in liquid nitrogen and then homogenized in 0.01 M PBS (pH 7.2), followed by centrifugation for 20 min at 2,000 g. The supernatant was used to determine the ATP, ATPase, alternative oxidase (AOX) and poly ADP-ribose polymerase (PARP) content via the ELISA method with an assay kit per the manufacturer’s instructions (Shanghai Enzyme-Linked Biotechnology Co., Ltd., China).

### Measurements of antioxidant enzyme activities

Frozen pistils (0.01 g) were ground into a fine powder in liquid nitrogen and then homogenized in 50 mM phosphate buffer (pH 7.0). The homogenate was centrifuged at 13,000 g for 15 min at 4°C, and the supernatant was stored in aliquots at −20°C until further analysis. Superoxide dismutase (SOD) activity was determined using the method of Giannopolitis and Ries (1977). The method of Maehly & Chance (1954) was used to determine the peroxidase (POD) activity, in which the rate of conversion of guaiacol to tetra-guaiacol was monitored at 470 nm. The activity of catalase (CAT) was determined using the method of Aebi (1983) with modifications, as described by Zhang et al. (2016). The ascorbate peroxidase (APX) activity was determined using the method of Bonnecarrère et al. (2011).

### Quantitative real-time PCR

Total RNA was extracted using TriPure Reagent (Aidlab Biotechnologies, Beijing, China) after triturating the tissue in liquid nitrogen, and the RNA concentration and quality were detected using a NanoDrop 1000 spectrophotometer (NanoDrop Technologies, Wilmington, DE, USA). RNA was reverse-transcribed into single-stranded cDNA using the ReverTra Ace qPCR RT Master Mix (TOYOBO, Shanghai, China). The cDNA was used as the template for PCR amplification. SYBR Green I (TOYOBO) was used as the fluorescent dye, and the Thermal Cycler Dice Real-Time System II (TaKaRa Biotechnology, Dalian, China) was used for real-time fluorescent qPCR analysis. The primers were designed using PRIMER 5 software and are listed in Supplementary Table 1. qRT-PCR was performed per the method of Feng et al. (2013), and the relative expression levels of the genes were analyzed using the 2^−ΔΔCT^ method.

### Statistical analysis

Data were processed using SPSS software 11.5 (IBM Corp., Armonk, NY, USA). The mean values and standard errors in the figures are from three replicates of independent experiments, unless otherwise stated. Student’s *t*-tests were conducted for normally distributed data. One-way analysis of variance (ANOVA) to compare the difference with a least significant difference test (LSD) at P ≤ 0.05 for the data within one genotype was conducted.

## Acknowledgments

We thank Prof. Jie Xiong from College of Life Sciences and Medicine of Zhejiang Sci-Tech University for useful discussion and experiment assistance.

## Author Contributions

GF and LT designed the research. HL, BF, JL and WF performed the experiments. GF and HL analyzed the data. HL and GF wrote the paper with contributions from all the authors. WW, TC and LL assisted in the determination of experimental data. SP and ZW refined the experimental design and provided advice. All authors read and approved the final manuscript.

## Figure legends

**Supplemental Figure 1** Effect of low-light stress at anthesis on dry matter weight of various parts of rice plants.

**Supplemental Figure 2** Effect of low-light stress on net photosynthesis in rice at anthesis.

**Supplemental Figure 3** Effects of low-light stress on rice pollen viability in rice at anthesis.

**Supplemental Figure 4** Effects of low-light stress on anther dehiscence in rice at anthesis.

**Supplemental Figure 5** Effects of low-light stress on PARP activity in pistil of rice at anthesis.

**Supplemental Figure 6** Effects of low-light stress on the expression of *HSPs* in pistil of rice at anthesis.

**Supplemental Figure 7** Effect of low-light stress on sugar signaling in pistil of rice at anthesis.

**Supplemental Table 1** Primer sequences used in quantitative Real-Time reverse transcription PCR

## References

Aebi H (1983) Catalase. In: H. U. Bergmeyer, ed. Methods of Enzymatic Analysis. Academic Press, Inc, New York, NY, USA. pp. 273–288

Amthor JS, Bar-Even A, Hanson AD, Millar AH, Stitt M, Sweetlove LJ, Tyerman SD (2019) Engineering strategies to boost crop productivity by cutting respiratory carbon loss. Plant Cell 31(2): 297–314

Baena-González E, Hanson J (2017) Shaping plant development through the SnRK1-TOR metabolic regulators. Curr Opin Plant Biol 35: 152–157

Barratt DH, Derbyshire P, Findlay K, Pike M, Wellner N, Lunn J, Feil R, Simpson C, Maule AJ, Smith AM (2009) Normal growth of Arabidopsis requires cytosolic invertase but not sucrose synthase. Proc Natl Acad Sci USA 106(31): 13124–13129

Bonnecarrère V, Borsani O, Díaz P, Capdevielle F, Blanco P, Monza J (2011) Response to photoxidative stress induced by cold in japonica rice is genotype dependent. Plant Sci 180(5): 726–732

Brennan T, Frenkel C (1977) Involvement of hydrogen peroxide in the regulation of senescence in pear. Plant Physiol 59(3): 411–416

Cárdenas L, McKenna ST, Kunkel JG, Hepler PK (2006) NAD(P)H oscillates in pollen tubes and is correlated with tip growth. Plant Physiol 142(4): 1460–1468

Certal AC, Almeida RB, Carvalho LM, Wong E, Moreno N, Michard E, Carneiro J, Rodriguéz-Léon J, Wu HM, Cheung AY, et al. (2008) Exclusion of a proton ATPase from the apical membrane is associated with cell polarity and tip growth in Nicotiana tabacum pollen tubes. Plant Cell 20(3): 614–634

Chen CT, Setter TL (2003) Response of potato tuber cell division and growth to shade and elevated CO_2_. Ann Bot 91(3): 373–381

Chen G, Chen H, Shi K, Raza MA, Bawa G, Sun X, Pu T, Yong T, Liu W, Liu J, et al. (2020) Heterogeneous Light Conditions Reduce the Assimilate Translocation Towards Maize Ears. Plants (Basel) 9(8): 987

Chen H, Li QP, Zeng YL, Deng F, Ren WJ (2019) Effect of different shading materials on grain yield and quality of rice. Sci Rep 9(1): 9992

Chen Q, Hu T, Li X, Song CP, Zhu JK, Chen L, Zhao Y (2022a) Phosphorylation of SWEET sucrose transporters regulates plant root:shoot ratio under drought. Nat Plants 8(1): 68–77

Chen Q, Zhang J, Li G (2022b) Dynamic epigenetic modifications in plant sugar signal transduction. Trends Plant Sci 27(4): 379–390

Colaneri AC, Tunc-Ozdemir M, Huang JP, Jones AM (2014) Growth attenuation under saline stress is mediated by the heterotrimeric G protein complex. BMC Plant Biol 14: 129

De Block M, Van Lijsebettens M (2011) Energy efficiency and energy homeostasis as genetic and epigenetic components of plant performance and crop productivity. Curr Opin Plant Biol 14(3): 275–282

Deng F, Wang L, Pu SL, Mei XF, Li SX, Li QP, Ren WJ (2018) Shading stress increases chalkiness by postponing caryopsis development and disturbing starch characteristics of rice grains. Agric For Meteorol 263: 49–58

Deng F, Zeng Y, Li Q, He C, Li B, Zhu Y, Zhou X, Yang F, Zhong X, Wang L (2021) Decreased anther dehiscence contributes to a lower fertilization rate of rice subjected to shading stress. Field Crop Res 273: 108291

Dhindsa RS, Plumb-Dhindsa P, Thorpe TA (1981) Leaf senescence: correlated with increased levels of membrane permeability and lipid peroxidation, and decreased levels of superoxide dismutase and catalase. J Exp Bot 32(1): 93–101

Dresselhaus T, Sprunck S, Wessel GM (2016) Fertilization Mechanisms in Flowering Plants. Curr Biol 26(3): R125–R139

Dröge-Laser W, Weiste C (2018) The C/S1 bZIP Network: A Regulatory Hub Orchestrating Plant Energy Homeostasis. Trends Plant Sci 23(5): 422–433

Dubois M, Gilles K, Hamilton J, Rebers P, Smith F (1956) Colorimetric method for determination of sugars and related substances. Anal Chem 28: 350–356

Feng BH, Yang Y, Shi YF, Shen HC, Wang HM, Huang QN, Xu X, Lü XG, Wu JL (2013) Characterization and genetic analysis of a novel rice spotted-leaf mutant HM_47_ with broad-spectrum resistance to Xanthomonas oryzae pv. Oryzae. J Integr Plant Biol 55(5): 473–483

Fernández-Milmanda GL, Ballaré CL (2021) Shade avoidance: expanding the color and hormone palette. Trends Plant Sci 26(5): 509–523

Ferrero-Serrano Á, Assmann SM (2016) The α-subunit of the rice heterotrimeric G protein, RGA1, regulates drought tolerance during the vegetative phase in the dwarf rice mutant d1. J Exp Bot 67(11): 3433–43

Ferrero-Serrano Á, Su Z, Assmann SM (2018) Illuminating the role of the Gα heterotrimeric G protein subunit, RGA1, in regulating photoprotection and photoavoidance in rice. Plant Cell Environ 41(2): 451–468

Fu G, Feng B, Zhang C, Yang Y, Yang X, Chen T, Zhao X, Zhang X, Jin Q, Tao L (2016) Heat stress is more damaging to superior spikelets than inferiors of rice (*Oryza sativa* L.) due to their different organ temperatures. Front Plant Sci 7: 1637

Fukuda N, Fujita M, Ohta Y, Sase S, Nishimura S, Ezura H (2008) Directional blue light irradiation triggers epidermal cell elongation of abaxial side resulting in inhibition of leaf epinasty in geranium under red light condition. Sci Hortic 115: 176–182

Fünfgeld MMFF, Wang W, Ishihara H, Arrivault S, Feil R, Smith AM, Stitt M, Lunn JE, Niittylä T (2022) Sucrose synthases are not involved in starch synthesis in Arabidopsis leaves. Nat Plants 8(5): 574–582

Ganguly S, Saha S, Vangaru S, Purkayastha S, Das D, Saha AK, Roy A, Das S, Bhattacharyya PK, Mukherjee S, et al. (2020) Identification and analysis of low light tolerant rice genotypes in field conditions and their SSR-based diversity in various abiotic stress tolerant lines. J Genet 99: 24

Ge Z, Bergonci T, Zhao Y, Zou Y, Du S, Liu MC, Luo X, Ruan H, García-Valencia LE, Zhong S, et al. (2017) Arabidopsis pollen tube integrity and sperm release are regulated by RALF-mediated signaling. Science 358(6370): 1596–1600

Ge Z, Cheung AY, Qu LJ (2019) Pollen tube integrity regulation in flowering plants: Insights from molecular assemblies on the pollen tube surface. New Phytol 222(2): 687–693

Giannopolitis CN, Ries SK (1977) Superoxide dismutases: I. Occurrence in higher plants. Plant Physiol 59: 309–314

Gommers CM, Visser EJ, St Onge KR, Voesenek LA, Pierik R (2013) Shade tolerance: when growing tall is not an option. Trends Plant Sci 18(2): 65–71

Goetz M, Guivarćh A, Hirsche J, Bauerfeind MA, González MC, Hyun TK, Eom SH, Chriqui D, Engelke T, Großkinsky DK, et al. (2017) Metabolic control of tobacco pollination by sugars and invertases. Plant Physiol 173(2): 984–997

Hackel A, Schauer N, Carrari F, Fernie AR, Grimm B, Kühn C (2006) Sucrose transporter *LeSUT1* and *LeSUT2* inhibition affects tomato fruit development in different ways. Plant J 45(2): 180–192

Hao LH, Wang WX, Chen C, Wang YF, Liu T, Li X, Shang ZL (2012) Extracellular ATP promotes stomatal opening of Arabidopsis thaliana through heterotrimeric G protein α subunit and reactive oxygen species. Mol Plant 5(4): 852–64

Hoffmann RD, Portes MT, Olsen LI, Damineli DSC, Hayashi M, Nunes CO, Pedersen JT, Lima PT, Campos C, Feijó JA, et al. (2020) Plasma membrane H^+^-ATPases sustain pollen tube growth and fertilization. Nat Commun 11(1): 2395

Horie T (2019) Global warming and rice production in Asia: Modeling, impact prediction and adaptation. Proc Jpn Acad Ser B Phys Biol Sci 95(6): 211–245

Huang JP, Tunc-Ozdemir M, Chang Y, Jones AM (2015) Cooperative control between AtRGS1 and AtHXK1 in a WD40-repeat protein pathway in Arabidopsis thaliana. Front Plant Sci 6: 851

Ishikawa A, Tsubouchi H, Iwasaki Y, Asahi T (1995) Molecular cloning and characterization of a cDNA for the alpha subunit of a G protein from rice. Plant Cell Physiol 36(2): 353–359

Islam M Rezaul, Feng B, Chen T, Fu W, Zhang C, Tao L, Fu G (2019) Abscisic acid prevents pollen abortion under high temperature stress by mediating sugar metabolism in rice spikelets. Physiol Plant 165(3): 644–663

Jagadish S (2020) Heat stress during flowering in cereals - effects and adaptation strategies. The New Phytol 226(6): 1567–1572

Jangam AP, Pathak RR, Raghuram N (2016) Microarray analysis of rice d1 (RGA1) mutant reveals the potential role of G-Protein alpha subunit in regulating multiple abiotic stresses such as drought, salinity, heat, and cold. Front Plant Sci 7: 11

Jiang N, Yu P, Fu W, Li G, Feng B, Chen T, Li H, Tao L, Fu G (2020) Acid invertase confers heat tolerance in rice plants by maintaining energy homoeostasis of spikelets. Plant Cell Environ 43(5): 1273–1287

Kan Y, Mu XR, Zhang H, Gao J, Shan JX, Ye WW, Lin HX (2022) TT2 controls rice thermotolerance through SCT1-dependent alteration of wax biosynthesis. Nat Plants 8(1): 53–67

Kim YJ, Kim MH, Hong WJ, Moon S, Kim ST, Park SK, Jung KH (2021) *OsMTD2*-mediated reactive oxygen species (ROS) balance is essential for intact pollen-tube elongation in rice. Plant J 107(4): 1131–1147

Kim YM, Heinzel N, Giese JO, Koeber J, Melzer M, Rutten T, Von Wirén N, Sonnewald U, Hajirezaei MR (2013) A dual role of tobacco hexokinase 1 in primary metabolism and sugar sensing. Plant Cell Environ 36(7): 1311–27

Kobata T, Sugawara M, Takatu S (2000) Shading during the early grain filling period does not affect potential grain dry matter increase in Rice. Agron J 92: 411–417

Kobata T, Yoshida H, Masiko U, Honda T (2013) Spikelet sterility is associated with a lack of assimilate in high-spikelet-number rice. Agron J 105: 1821–1831

Kono M, Kawaguchi H, Mizusawa N, Yamori W, Suzuki Y, Terashima I (2020) Far-Red Light Accelerates Photosynthesis in the Low-Light Phases of Fluctuating Light. Plant Cell Physiol 61(1): 192–202

Kumar A, Panda D, Mohanty S, Biswal M, Dey P, Dash M, Sah RP, Kumar S, Baig MJ, Behera L (2020) Role of sedoheptulose-1,7 bisphosphatase in low light tolerance of rice (*Oryza sativa* L.). Physiol Mol Biol Plants 26(12): 2465– 2485

Lee CP, Millar AH (2016) The plant mitochondrial transportome: Balancing metabolic demands with energetic constraints. Trends Plant Sci 21(8): 662–676

Li DD, Guan H, Li F, Liu CZ, Dong YX, Zhang XS, Gao XQ (2017) Arabidopsis shaker pollen inward K^+^ channel SPIK functions in SnRK1 complex-regulated pollen hydration on the stigma. J Integr Plant Biol 59(9): 604–611

Li HJ, Yang WC (2018) Ligands switch model for pollen-tube integrity and burst. Trends Plant Sci 23(5): 369–372

Li G, Chen T, Feng B, Peng S, Tao L, Fu G (2021) Respiration, rather than photosynthesis, determines rice yield loss under moderate high-temperature conditions. Front Plant Sci 12: 678653

Li Y, Guan K, Schnitkey GD, DeLucia E, Peng B (2019) Excessive rainfall leads to maize yield loss of a comparable magnitude to extreme drought in the United States. Glob Chang Biol 25(7): 2325–2337

Long SP, Taylor SH, Burgess SJ, Carmo-Silva E, Lawson T, De Souza AP, Leonelli L, Wang Y (2022) Into the shadows and back into sunlight: photosynthesis in fluctuating light. Annu Rev Plant Biol 73: 617–648

Maehly AC, Chance B (1954) The assay of catalases and peroxidases. Methods Biochem Anal 1: 357–424

Margalha L, Confraria A, Baena-González E (2019) SnRK1 and TOR: modulating growth-defense trade-offs in plant stress responses. J Exp Bot 70(8): 2261–2274

Ma Y, Dai X, Xu Y, Luo W, Zheng X, Zeng D, Pan Y, Lin X, Liu H, Zhang D, et al. (2015) COLD1 confers chilling tolerance in rice. Cell 160(6): 1209–1221

Misra S, Wu Y, Venkataraman G, Sopory SK, Tuteja N (2007) Heterotrimeric G-protein complex and G-protein-coupled receptor from a legume (Pisum sativum): role in salinity and heat stress and cross-talk with phospholipase C. Plant J 51(4): 656–669

O’Leary BM, Oh GGK, Lee CP, Millar AH (2020) Metabolite regulatory interactions control plant respiratory metabolism via target of rapamycin (TOR) kinase activation. Plant Cell 32: 666–682

Olearczyk JJ, Stephenson AH, Lonigro AJ, Sprague RS (2004a) Heterotrimeric G protein Gi is involved in a signal transduction pathway for ATP release from erythrocytes. Am J Physiol Heart Circ Physiol 286(3): H940–H945

Olearczyk JJ, Stephenson AH, Lonigro AJ, Sprague RS (2004b) NO inhibits signal transduction pathway for ATP release from erythrocytes via its action on heterotrimeric G protein Gi. Am J Physiol Heart Circ Physiol 287(2): H748– H754

Oliveira PF, Lopes IA, Barrias C, Rebelo da Costa AM (2004) H^+^-ATPase of crude homogenate of the outer mantle epithelium of Anodonta cygnea. Comp Biochem Physiol A Mol Integr Physiol 139(4): 425–432

Peng Y, Chen L, Li S, Zhang Y, Xu R, Liu Z, Liu W, Kong J, Huang X, Wang Y, et al. (2018) BRI1 and BAK1 interact with G proteins and regulate sugar-responsive growth and development in Arabidopsis. Nat Commun 9(1): 1522

Peskan-Berghöfer T, Neuwirth J, Kusnetsov V, Oelmüller R (2005) Suppression of heterotrimeric G-protein beta-subunit affects anther shape, pollen development and inflorescence architecture in tobacco. Planta 220(5): 737–746

Plaxton WC, Podestá FE (2006) The functional organization and control of plant respiration. Crit Rev Plant 25(2): 159–198

Proctor J (2021) Atmospheric opacity has a nonlinear effect on global crop yields. Nat Food 2(3): 166–173

Roitsch T, Gonzalez MC (2004) Function and regulation of plant invertases: Sweet sensations. Trends Plant Sci 9(12): 606–613

Rottmann T, Zierer W, Subert C, Sauer N, Stadler R (2016) STP10 encodes a high-affinity monosaccharide transporter and is induced under low-glucose conditions in pollen tubes of Arabidopsis. J Exp Bot 67(8): 2387–2399

Ruan YL (2014) Sucrose metabolism: gateway to diverse carbon use and sugar signaling. Annu Rev Plant Biol 65: 33–67

Schwenkert S, Fernie AR, Geigenberger P, Leister D, Möhlmann T, Naranjo B, Neuhaus HE (2022) Chloroplasts are key players to cope with light and temperature stress. Trends Plant Sci 27(6): 577–587

Shao L, Li G, Zhao Q, Li Y, Sun Y, Wang W, Cai C, Chen W, Liu R, Luo W, et al. (2020) The fertilization effect of global dimming on crop yields is not attributed to an improved light interception. Glob Chang Biol 26(3): 1697–1713

Shao L, Liu Z, Li H, Zhang Y, Dong M, Guo X, Zhang H, Huang B, Ni R, Li G, et al. (2021) The impact of global dimming on crop yields is determined by the source-sink imbalance of carbon during grain filling. Glob Chang Biol 27(3): 689–708

Slattery RA, Walker BJ, Weber APM, Ort DR (2018) The impacts of fluctuating light on crop performance. Plant Physiol 176(2): 990–1003

Sosso D, Luo D, Li QB, Sasse J, Yang J, Gendrot G, Suzuki M, Koch KE, McCarty DR, Chourey PS, et al. (2015) Seed filling in domesticated maize and rice depends on SWEET-mediated hexose transport. Nat Genet 47(12): 1489–1493

Stein O, Granot D (2019) An Overview of Sucrose Synthases in Plants. Front Plant Sci 10: 95

Tollenaar M, Fridgen J, Tyagi P, Stackhouse PWJr, Kumudini S (2017) The contribution of solar brightening to the US maize yield trend. Nat Clim Change 7: 275

Urano D, Colaneri A, Jones AM (2014) Gα modulates salt-induced cellular senescence and cell division in rice and maize. J Exp Bot 65(22): 6553–6561

Wang F, Sanz A, Brenner ML, Smith A (1993) Sucrose synthase, starch accumulation, and tomato fruit sink strength. Plant Physiol 101(1): 321–327

Wang J, Wang A, Luo Q, Hu Z, Ma Q, Li Y, Lin T, Liang X, Yu J, Foyer CH, Shi K (2022) Glucose sensing by regulator of G protein signaling 1 (RGS1) plays a crucial role in coordinating defense in response to environmental variation in tomato. New Phytol 10.1111/nph.18356

Wang L, Deng F, Ren WJ (2015) Shading tolerance in rice is related to better light harvesting and use efficiency and grain filling rate during grain filling period. Field Crop Res 180: 54– 62

Wu Y, Xu X, Li S, Liu T, Ma L, Shang Z (2007) Heterotrimeric G-protein participation in Arabidopsis pollen germination through modulation of a plasmamembrane hyperpolarization-activated Ca^2+^-permeable channel. New Phytol 176(3): 550–559

Xiang X, Zhang P, Yu P, Zhang Y, Yang Z, Sun L, Wu W, Khan RM, Abbas A, Cheng S, et al. (2019) LSSR1 facilitates seed setting rate by promoting fertilization in rice. Rice 12(1): 31

Xu Y, Yang J, Wang Y, Wang J, Yu Y, Long Y, Wang Y, Zhang H, Ren Y, Chen J, et al. (2017) *OsCNGC13* promotes seed-setting rate by facilitating pollen tube growth in stylar tissues. PLoS Genet 13(7): e1006906

Yang J, Luo D, Yang B, Frommer WB, Eom JS (2018) SWEET11 and 15 as key players in seed filling in rice. New Phytol 218(2): 604–615

Yu P, Jiang N, Fu W, Zheng G, Li G, Feng B, Chen T, Ma J, Li H, Tao L, et al. (2020) ATP hydrolysis determines cold tolerance by regulating available energy for glutathione synthesis in rice seedling plants. Rice 13(1): 23

Yu SM, Lo SF, Ho TD (2015) Source-sink communication: Regulated by hormone, nutrient, and stress cross-signaling. Trends Plant Sci 20(12): 844–857

Yu X, Zhang X, Zhao P, Peng X, Chen H, Bleckmann A, Bazhenova A, Shi C, Dresselhaus T, Sun MX (2021) Fertilized egg cells secrete endopeptidases to avoid polytubey. Nature 592(7854): 433–437

Zhang CX, Fu GF, Yang XQ, Yang YJ, Zhao X, Chen TT, Zhang XF, Jin QY, Tao LX (2016) Heat stress effects are stronger on spikelets than on flag leaves in rice due to differences in dissipation capacity. J Agron C Sci 202(5): 394–408

Zhang CX, Li GY, Chen TT, Feng BH, Fu WM, Yan JX, Islam MR, Jin QY, Tao LX, Fu GF (2018a) Heat stress induces spikelet sterility in rice at anthesis through inhibition of pollen tube elongation interfering with auxin homeostasis in pollinated pistils. Rice 11(1): 14

Zhang CX, Feng BH, Chen TT, Fu WM, Li HB, Li GY, Jin QY, Tao LX, Fu GF (2018b) Heat stress-reduced kernel weight in rice at anthesis is associated with impaired source-sink relationship and sugars allocation. Environ Exp Bot 155: 718–733

Zhang H, Xie P, Xu X, Xie Q, Yu F (2021) Heterotrimeric G protein signalling in plant biotic and abiotic stress response. Plant Biol (Stuttg) 1: 20–30

Zhang J, Huang Q, Zhong S, Bleckmann A, Huang J, Guo X, Lin Q, Gu H, Dong J, Dresselhaus T, et al. (2017) Sperm cells are passive cargo of the pollen tube in plant fertilization. Nat Plants 3: 17079

Zhang L, Fang K, Lin J (2005) Heterotrimeric G protein α-subunit is localized in the plasma membrane of Pinus bungeana pollen tubes. Plant Sci 169(6): 1066–1073

Zhang W, Jeon BW, Assmann SM (2011) Heterotrimeric G-protein regulation of ROS signalling and calcium currents in Arabidopsis guard cells. J Exp Bot 62(7): 2371–2379

Zhang YJ (1977) Determination of glucose, fructose, sucrose and starch in fruit and vegetable with anthrone colorimetric method. Chin J Anal Chem 5(3): 167–171

Zhong S, Li L, Wang Z, Ge Z, Li Q, Bleckmann A, Wang J, Song Z, Shi Y, Liu T, et al. (2022) RALF peptide signaling controls the polytubey block in Arabidopsis. Science 375(6578): 290–296

Zhou LZ, Qu LJ, Dresselhaus T (2021) Stigmatic ROS: regulator of compatible pollen tube perception?. Trends Plant Sci 26(10): 993–995

